# Structural modeling of cytokine-receptor-JAK2 signaling complexes using AlphaFold Multimer

**DOI:** 10.1101/2023.06.14.544971

**Authors:** Irina D. Pogozheva, Stanislav Cherepanov, Sang-Jun Park, Malini Raghavan, Wonpil Im, Andrei L. Lomize

**Affiliations:** Department of Medicinal Chemistry, College of Pharmacy, University of Michigan, Ann Arbor, MI 48109, United States; Biophysics Program, University of Michigan, Ann Arbor, MI 48109, United States; Departments of Biological Sciences and Chemistry, Lehigh University, Bethlehem, PA 18015, United States; Department of Microbiology and Immunology, University of Michigan Medical School, Ann Arbor, MI 48109, United States

**Keywords:** granulocyte colony-stimulating factor 3 receptor, growth hormone receptor, erythropoietin receptor, prolactin receptor, thrombopoietin receptor

## Abstract

Homodimeric class 1 cytokine receptors include the erythropoietin (EPOR), thrombopoietin (TPOR), granulocyte colony-stimulating factor 3 (CSF3R), growth hormone (GHR), and prolactin receptors (PRLR). They are cell-surface single-pass transmembrane (TM) glycoproteins that regulate cell growth, proliferation, and differentiation and induce oncogenesis. An active TM signaling complex consists of a receptor homodimer, one or two ligands bound to the receptor extracellular domains and two molecules of Janus Kinase 2 (JAK2) constitutively associated with the receptor intracellular domains. Although crystal structures of soluble extracellular domains with ligands have been obtained for all the receptors except TPOR, little is known about the structure and dynamics of the complete TM complexes that activate the downstream JAK-STAT signaling pathway. Three-dimensional models of five human receptor complexes with cytokines and JAK2 were generated using AlphaFold Multimer. Given the large size of the complexes (from 3220 to 4074 residues), the modeling required a stepwise assembly from smaller parts with selection and validation of the models through comparisons with published experimental data. The modeling of active and inactive complexes supports a general activation mechanism that involves ligand binding to a monomeric receptor followed by receptor dimerization and rotational movement of the receptor TM α-helices causing proximity, dimerization, and activation of associated JAK2 subunits. The binding mode of two eltrombopag molecules to TM α-helices of the active TPOR dimer was proposed. The models also help elucidating the molecular basis of oncogenic mutations that may involve non-canonical activation route. Models equilibrated in explicit lipids of the plasma membrane are publicly available.

## Introduction

Cytokines are small secreted glycoproteins that regulate hematopoiesis, neurogenesis, adaptive and innate immunity, lactation, reproduction, growth, and metabolism through binding to cognate receptors [1]. Cytokine receptors are cell-surface glycoproteins with single transmembrane (TM) α-helix. They lack kinase activity and, therefore, rely on cytoplasmic tyrosine kinases to mediate intracellular processes. In particular, class 1 and 2 cytokines act *via* binding to cytokine receptors on the surface of target cells to activate the cytoplasmic non-receptor protein tyrosine kinase from the Janus Kinases (JAK) family that initiates the downstream JAK-STAT signaling pathway. The cytokine-initiated signaling involves five consecutive steps: (1) binding of a cytokine to a specific receptor and formation of the active receptor dimer; (2) activation of the receptor-associated JAKs by dimerization and *trans*-phosphorylation in their activation loops; (3) phosphorylation of receptor tyrosine residues by the activated JAK; (4) binding of a STAT transcription factor to phosphotyrosines of receptor, leading to phosphorylation, dimer rearrangement, and nuclear translocation of STAT to drive the expression of cytokine-responsive genes; and (5) switching off the activated receptor by tyrosine phosphatases (SHPs), Suppressors Of Cytokine Signaling (SOCS), receptor internalization and downregulation [2]. Human genomes encode more than 50 cytokine receptors, four members of JAK family (JAK1, JAK2, JAK3, and TYK2), seven STATs (STAT1-4, STAT5a, STAT5b, and STAT6), two SHPs (SHP1 and SHP2), and eight SOCS (SOCS1-7 and CIS) [1–3]. Specific members of the JAK, STAT and SOCS family are linked to individual receptors (**Table S1**).

Cytokines of the JAK-STAT pathway are α-helical proteins that form either 4-α-helical bundles with up-up-down-down topology (class 1, **Figure S1)** or structures with 5-6 antiparallel α-helices arranged in an up-down fashion (class 2). The class 1 cytokine receptors are the largest group of 34 proteins encoded by human genome [4]. These single-pass TM proteins have different lengths, domain architectures, and quaternary structures. The class 1 receptor family includes five subfamilies: (1) homodimeric receptors that bind one or two ligands per receptor pair; (2, 3 and 4) subfamilies of interleukin (IL) receptor: the IL-12/23, IL-2, and IL-6 receptors forming heterodimers, heterotrimers or heterotetramers with ligand:receptor stoichiometries of 1:2, 1:3 and 2:4; and (5) a subfamily of IL-3 interleukin receptors forming a 12-meric complex composed of 8 receptors and 4 cytokine molecules [2].

The class I subfamily of homodimeric receptors includes receptors for erythropoietin (EPOR), growth hormone (GHR), prolactin (PRLR), thrombopoietin (TPOR, also called MPL or CD110), granulocyte colony-stimulating factor 3 (CSF3R), and leptin. These receptors (**Figure 1**) use two identical chains, each composed of the β-structural extracellular domain (ECD) responsible for ligand binding, a single-helical TM domain (TMD) driving receptor dimerization, and a disordered intracellular domain (ICD) responsible for JAK2 binding and STAT signaling [2]. EPOR, GHR, and PRLR are structurally simple [5–7] with an ECD containing a single cytokine homology module (CHM) formed by two fibronectin type III (FnIII) domains, D1 and D2. The membrane-distal D1 domain carries two conserved disulfides, whereas the membrane-proximal D2 domain features a characteristic WSXWS motif [8], replaced by the YGEFS motif in GHR (**Figure 1C**). The TPOR ECD is twice as large, as composed of four FnIII domains: D1 and D2 forming the CHM1, D3 and D4 forming the CHM2 [9, 10]. The ECD of the long-chain CSF3R contains six domains: the immunoglobulin-like (Ig-like) D1 domain, two FnIII domains, D2 and D3, forming CHM, and three extra FnIII domains, D4, D5, and D6 [11]. The long-chain leptin receptor with a more complex domain architecture [12] will not be studied here.

**Figure 1.**
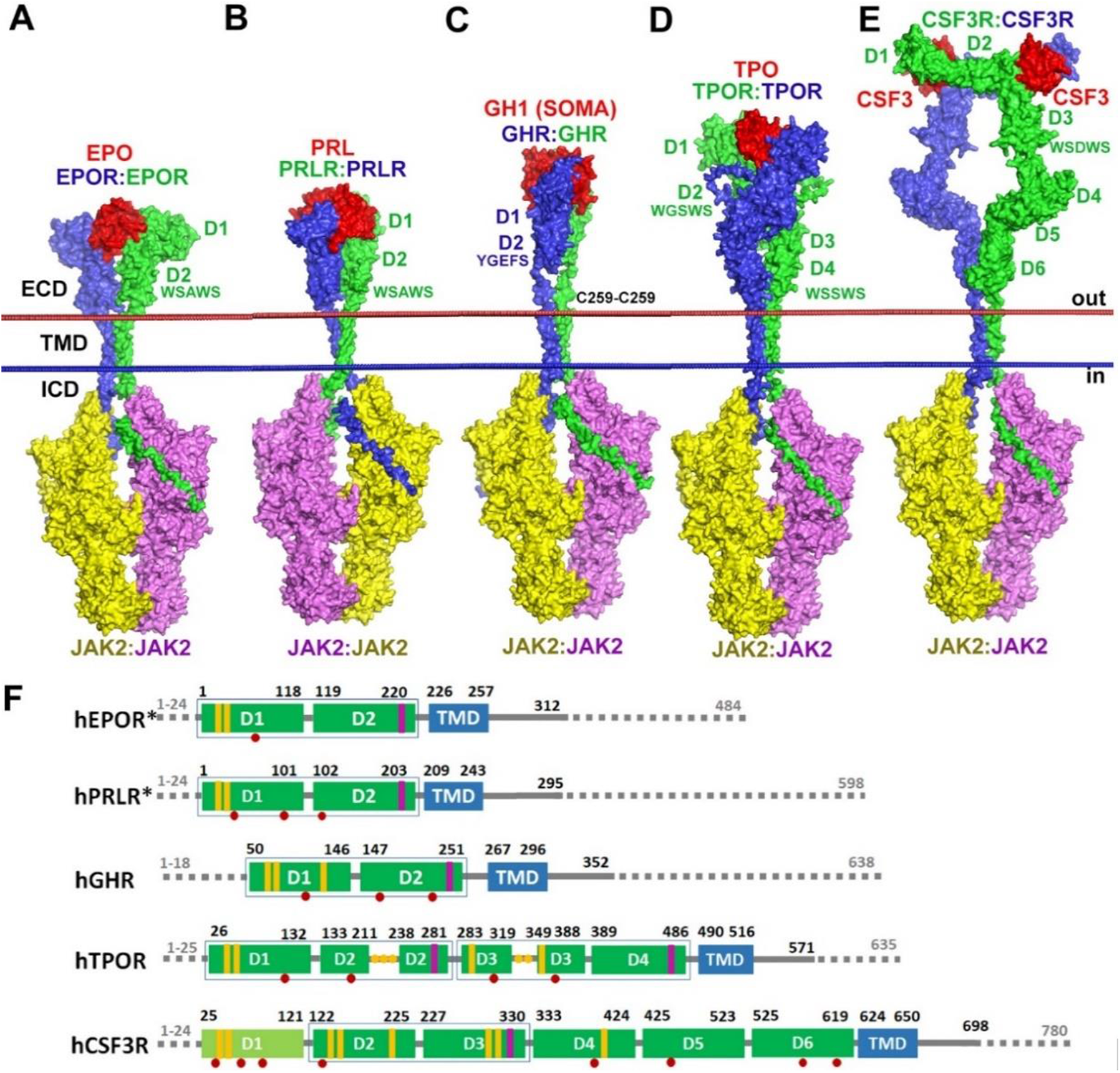
AlphaFold Multimer (AFM)-based models of active signaling complexes of human homodimeric class 1 cytokine receptors: EPOR (**A**), PRLR (**B**), GHR (**C**), TPOR (**D**), CSF3R (**E**). The complexes are composed of receptor homodimers, one (**A-D**) or two (**E**) ligands, and two JAK2 molecules bound to the intracellular domains (ICD) of receptors. Molecules are shown by surface representation and colored red for ligand, blue and green for receptor subunits, yellow and pink for JAK2 subunits. The extracellular domains (from D1 to D6) and WSXWS motifs of the receptors are indicated for each complex. The GHR complex has an intermolecular C259-C259 disulfide bond. Hydrophobic membrane boundaries are shown as red (extracellular side) and blue (intracellular side) spheres. (**F**) Domain architecture of the five cytokine receptors studied. The dark green boxes indicate Fibronectin typeIII (FnIII) domains. The ight green boxes indicate Immunoglobulin-like (Ig-like) domains. Blue boxes indicate TMDs. The boxes around two domains indicate cytokine homology modules (CHMs). The grey lines indicate unstructured regions or signal sequences. The yellow bars indicate disulfides. The purple bars indicate WSXWS motifs. The red circles indicate N-glycosylation sites. The, yellow circles indicate cysteine residues in loops of D2 and D3 domains of TPOR, which may form disulfides or metal-bound clusters. The dashed lines indicate disordered regions that have been omitted in the final models, but included during some of our AFM calculations. Asterisks indicate receptors with residue numbers corresponding to mature proteins lacking signal sequences.

Crystal structures of 1:2 complexes of natural cytokines with soluble receptor ECDs have been solved for human erythropoietin (EPO)-EPOR [13], human somatotropin (GH1)-GHR [14], and human prolactin (PRL) with rat PRLR [15] (PDB IDs: **1EER, 3HHR, 3NPZ**, respectively). In these crystal structures, two similar receptor chains create an interface for binding asymmetric surfaces of a cytokine molecule. Site 1 is formed by helices α1, α4 and the loop connecting α3 and α4, while site 2 is composed of α1 and α3 helices. Crystallographic and biophysical studies demonstrated that a cytokine initially binds to a single receptor *via* the high-affinity site 1 [16, 17] and then to the second receptor through the low-affinity site 2 [18]. The largest CSF3R-CSF3 complex has a different 2:2 receptor-ligand stoichiometry and represents the association of two 1:1 units [11]. The CSF3 binding site is formed by the CHM (D2 and D3) of one chain and the Ig-like domain (D1) of the other chain. At present, there are no experimental structures of TPOR or its domains. Hence, experimentally-based computational models have been proposed for the TPOR TM-ICD in complex with JAK2 dimer [19] or for the full-length human TPOR in complexes with thrombopoietin (TPO) or oncogenic calreticulin (CRT) mutants that bind to TPOR to cause its aberrant activation in myeloproliferative neoplasms (MPNs) [20, 21].

In the absence of experimental atomic-level structures of cytokine receptor signaling complexes, computational modeling provides a valuable alternative. A transformative breakthrough in the protein structure prediction has recently been achieved by developing a new deep learning AlphaFold method that uses co-evolutionary and structural information [22]. AlphaFold version 2.0 (AF2) produces models of nearly experimental quality for single-chain proteins and outperforms other methods in predicting contact interfaces of multi-domain proteins [23, 24] and protein complexes [25–27], including TM homo- and heterodimers [28, 29]. The high speed and quality of predictions by AF2 justified its applications on a proteomic scale [30]. More than 200 million protein models of single-chain proteins from 48 organisms were generated using this method and deposited into the AlphaFold DataBase [31]. The recently released AlphaFold Multimer (AFM) was recognized as a best computational tool for modeling protein complexes [32].

In this study we used the publicly available AFM ColabFold version [33] to model active signaling complexes of human class 1 homodimeric cytokine receptors, including EPOR, PRLR, GHR, TPOR and CSF3R. Each signaling complex is composed of one or two (for CSF3R) cytokines bound to a receptor homodimer interacting with JAK2 dimer. The accuracy of the models was verified through comparison with published experimental data. Comparison of ligand-free and cytokine-bound models revealed molecular mechanisms of receptor activation leading to the dimerization and activation of the receptor-bound JAK2. These models reveal atomic details of protein-protein interactions, demonstrating conformational changes and structural flexibilities in the ECDs, TMDs, and ICDs of receptors and JAK2 domains in the signaling complexes. This understanding aids in deciphering the nature of oncogenic mutations.

## Results

### Stepwise modeling with AFM

The direct modeling of complexes composed of 5 or 6 proteins using AFM was not feasible due to their very large size and multi-domain architecture. Therefore, for each cytokine receptor, we separately calculated several smaller overlapping parts and assembled them to a complete ligand-receptor-kinase complex. The modeling was performed in four steps (**Figure 2**).

**Figure 2.**
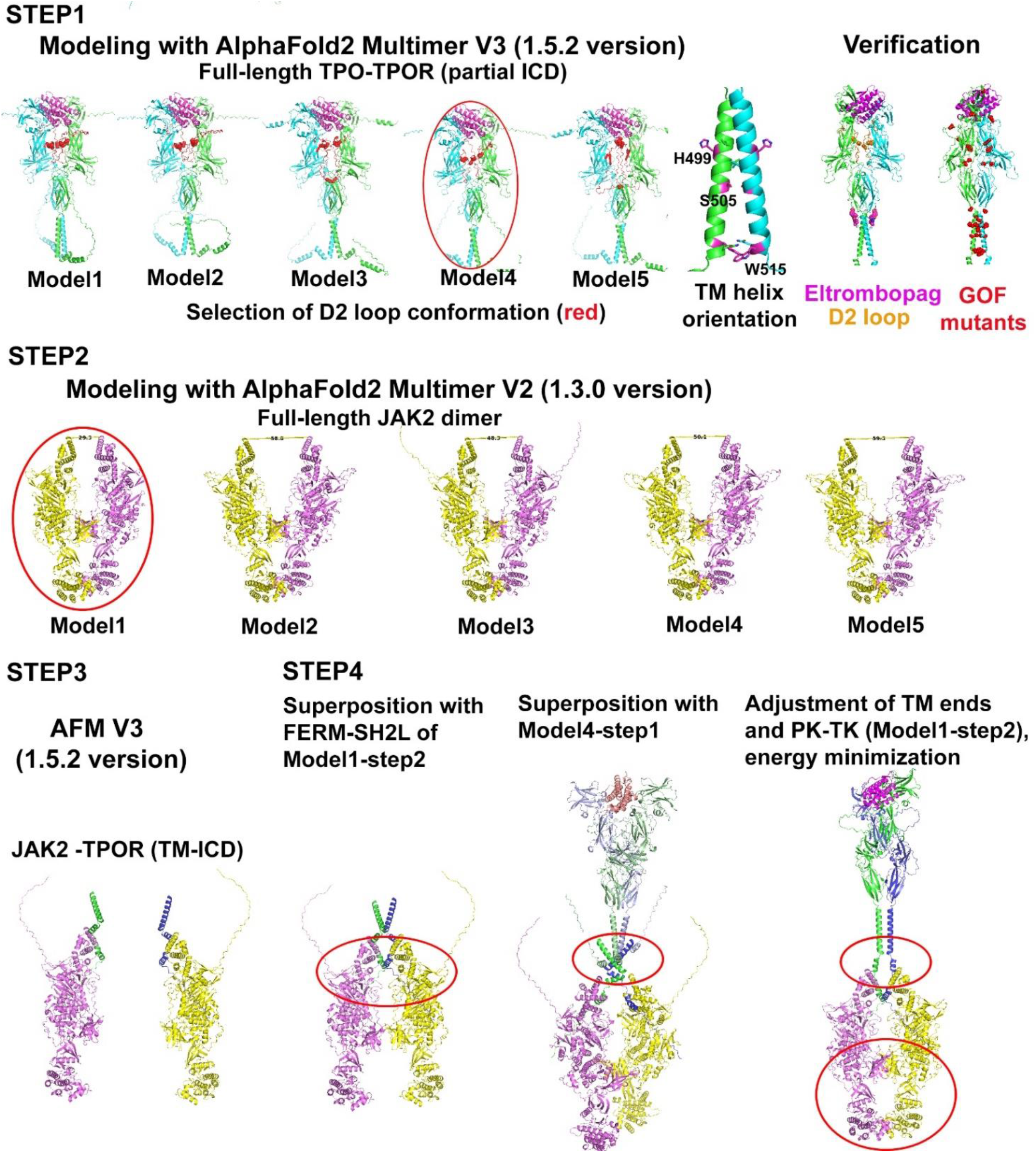
Four-step AMF-modeling of cytokine receptor signaling complexes (exemplified by hTPOR). <u>Step1</u>: Modeling with AFM V3 of TPO-TPOR active (1:2) complex. TPOR subunits are colored green and blue, TPO is colored purple. Five calculated models demonstrate similar conformations of ECDs except for the flexible D2 loop (residues 187-238 colored red) forming different intramolecular disulfides (shown as red spheres). The more frequent are the C193-C323, C194-C241, and C211-C322 disulfides. Four of five models demonstrate left-handed TM helical dimers with S505 at the dimerization interface (*a*-position of the heptad repeat motif) and H499 facing the lipid bilayer (*b*-position). Model 4 containing the most frequent disulfides and TMD helix arrangement was selected for the further calculations. Data supporting model 4 include: (1) the consistency of the TM α-helix arrangement with the *cc-TPOR-I* fusion construct between the dimeric coiled-coil of *Put3* and TMD of mTPOR that displays constitutive activity [53]; (2) the docking of two eltrombopag molecules (shown by purple spheres) to the TMD; and (3) the localization of the majority of GOF mutations (shown by red spheres) at flexible structural elements (D2 loop and TMDs) participating in dimerization interfaces. <u>Step2</u>: modeling of JAK2 dimers. JAK2 subunits are colored pink and yellow. AFM V2 and V3 models produce similar structures of JAK2 dimers with the PK-PK dimerization interface but different distances between FERM domains (Cα-Cα distances between two L244 varies between 30 and 60 Å). The model 1 with the minimal distance between FERM domains was selected for the further calculations. <u>Step3</u>. Modeling of the JAK2 monomer (colored pink or yellow) with TPOR TM α-helix and the ICD domain (colored blue and green). AFM V2 and V3 produced rather similar models. <u>Step4.</u> Structural superposition of two monomer models (from step 3) with FERM-SH2L domains of JAK2 dimer (model 1 from step2) followed by docking of the ligand-bound dimer (model 4 from step 1), adjustment of TM helix ends, substitution of PK and TK domains by those from JAK2 dimer (model 1 from step 2), and model refinement by energy minimization.

At the first step, we generated complexes of cytokines with their receptor homodimers that included ECDs, TMDs, and parts of ICDs (**Figure 1F**). In addition, we modeled dimers of peptides representing TM and juxtamembrane regions (TM-JM peptides) and dimers of ligand-free-receptors and compared them with the corresponding ligand-bound dimers. At the second step, we modeled the active dimer of human JAK2. At the third step, we produced complexes of a monomeric JAK2 with a TMD-ICD fragment of each receptor.

At the first three steps, we generated for each protein complex dozens of various structures using AFM V2 and V3 with different random seed numbers. Top score models varied in conformations of loops, orientations of TM α-helices, and, in some cases, in domain arrangement. Different sets of convergent models we compared with available crystal, NMR, and cryo-EM structures and validated though previously published mutagenesis, protein engineering, oncogenic mutations, and other experimental data to select one model the most compatible with these data. In case of multiple alternative structures and insufficient experimental data for an unambiguous model selection, we performed an additional modeling for protein sequences of various length, domain architecture, subunit stoichiometry, and for constitutively active mutants of TMDs.

At the fourth step, the full-length active signaling complex of each receptor was assembled from the best structures of ligand-bound receptor dimers and ICD-bound kinase dimers selected at the previous steps. The models of five receptor-cytokine signaling complexes were determined with rather high reliability scores for most structural domains, but lower reliability for loops and TM helices (**Figure S2**).

### Step 1: Modeling and validation of the receptor homodimers with and without ligands

#### Complexes of EPOR, GHR, and PRLR homodimers with cytokines

Models for complexes of cytokines with their cognate receptors were generated by AFM for five human receptors and for the extensively studied murine EPOR and validated using available experimental data (**Figure 2**). The models of dimeric short-chain receptors, EPOR, GHR, and PRLR, demonstrate similar structures of ECDs, but more diverse arrangement of TM helices.

The ECDs in the models of active dimers are in a good agreement with available experimental data. Indeed, ECDs of five cytokine-bound receptor dimers, EPO-EPOR_2_, GH1-GHR_2_, PRL-PRLR_2_, CSH1-PRLR_2_, and GH1-PRLR_2_, superimpose well with the corresponding crystal structures (PDB IDs: **1EER, 3HHR, 3NPZ, 1F6F,** and **1BP3,** respectively) with the Cα-atom root-mean-square deviations (RMSD) less than 2 Å (**Table S2**). Main residues involved in ligand-receptor interactions (**Figure 3**) are the same as in the corresponding experimental structures [13–16, 34]. For example, hydrophobic and aromatic residues, such as F93, F205, and M150 in hEPOR, W122 and W187 (W104 and W169 in mature protein) in hGHR, and W72 and W139 in hPRLR, contribute significantly to the hormone-receptor interactions.

**Figure 3.**
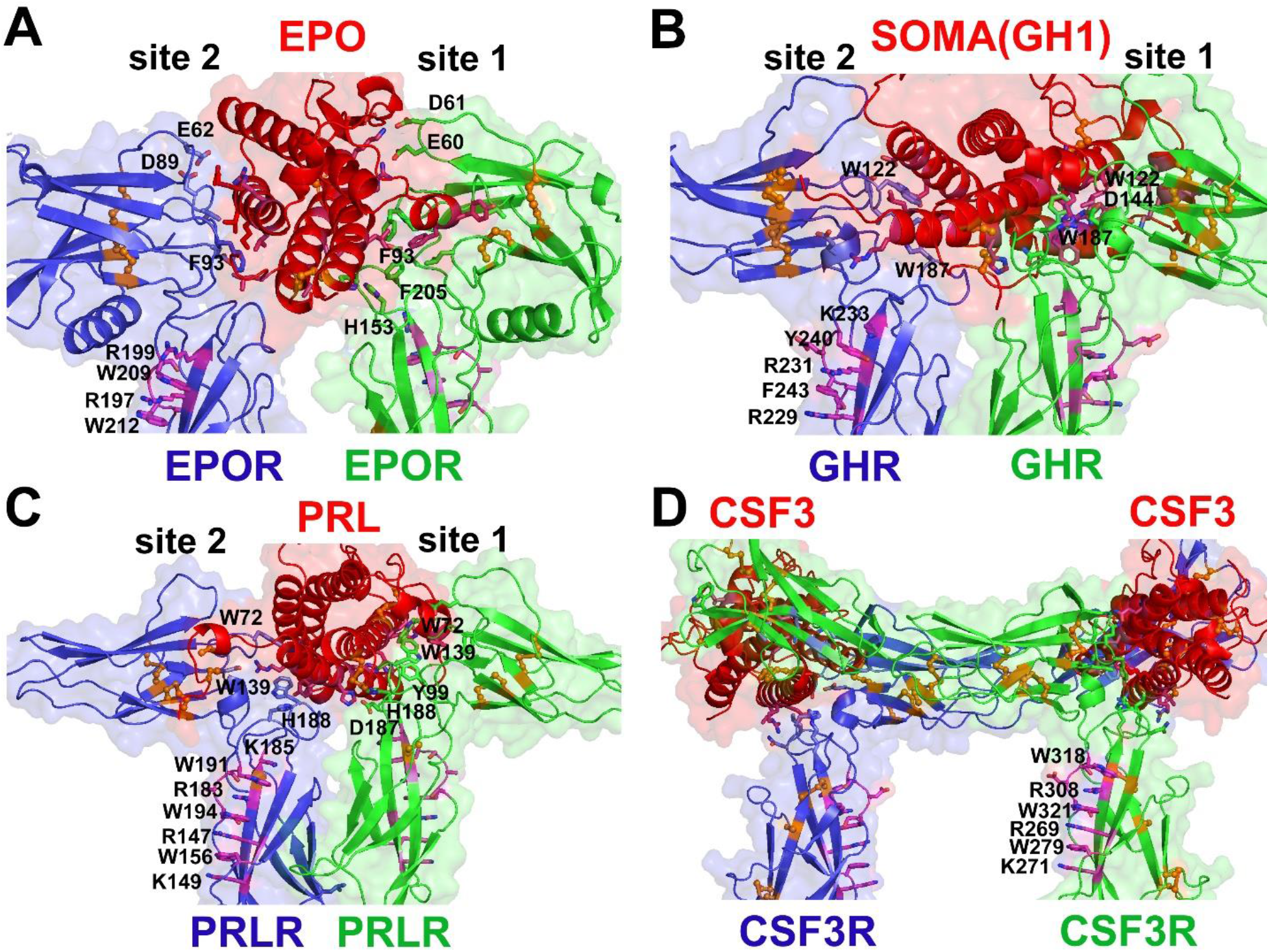
Ligand binding pockets in models of active homodimers of class 1 cytokine receptors: EPOR (**A**), GHR (**B**), PRLR (**C**), CSF3R (**D**) with bound cytokine ligands. Protein molecules are shown by semi-transparent surfaces and cartoon representations are colored red for ligands, and blue and green for receptor subunits. Interacting receptor and cytokine residues are shown as sticks. Cysteine residues are shown as balls-and-sticks colored orange. Residues involved in the WSXWS signature motif are shown as purple sticks. A set of interdigitated arginine and tryptophan residues from this motif together with neighboring tryptophan, arginine, and lysine residues participate in the network of cation-π interactions.

Models of ligand-bound EPOR, GHR, PRLR, and TPOR show a significant asymmetry of ECDs of two receptor chains. Binding of ligands with asymmetric binding surfaces (sites 1 and 2) induces movement of D1-D2 domains of both chains relative each other in vertical (along the membrane normal, *z*-axis) and horizontal (in the *xy* plane) directions. The vertical shift (by 5 to 8 Å) of the site 1 ECD is translated to the upward piston movement of the corresponding TM α-helix. Thus, TM helices of the dimer are positioned in the membrane at different heights (**Figure 4**). The vertical shift is pronounced in models of the short-chain receptors, but is not seen in models of multi-domain long-chain receptors. The asymmetry is absent in complexes of hEPOR dimers with two similar molecules of peptide mimetics (PDB IDs: **1EBA, 1EBP**) and in the ligand-free receptor dimers (see below).

**Figure 4.**
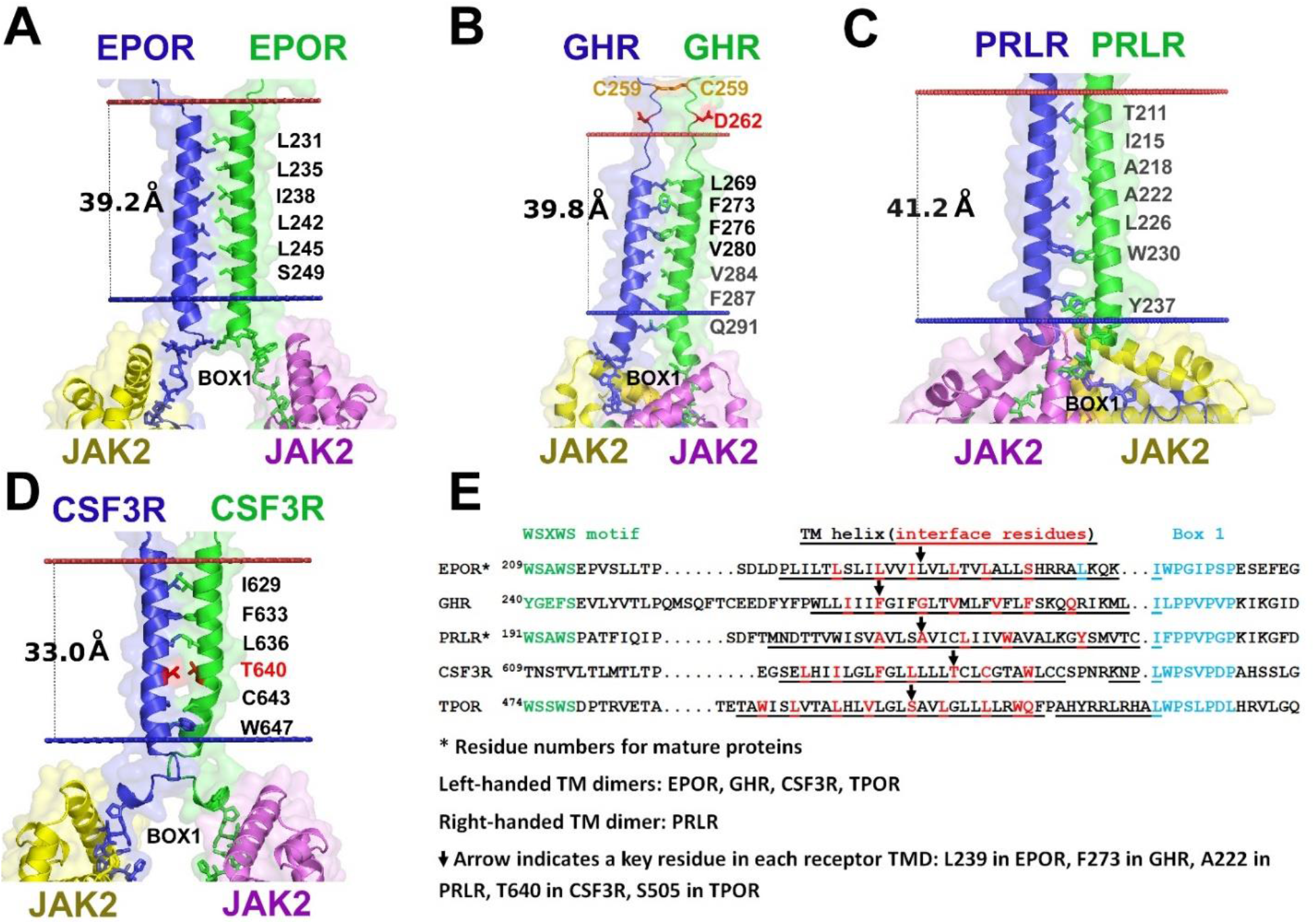
The TM α-helical dimers with predicted locations of membrane boundaries in AFM models of signaling complexes of EPOR (**A**), GHR (**B**), PRLR (**C**) and CSF3R (**D**). Each complex is composed of two receptor molecules (colored blue and green), bound cytokine(s) (not shown) and subunits of a JAK2 homodimer (colored yellow and pink). The TM α-helices form left-handed dimers with positive crossing angles (*via* extended leucine zipper heptad repeat pattern) for EPOR, GHR and CSF3R, and a right-handed dimer for PRLR (*via* AxxxA^222^xxxL motif). Residues from the TMD dimerization interface and Box1 residues are shown as sticks. Cysteine residues are shown as balls-and-sticks colored orange, and C259 residues of the intermolecular disulfide in the active complex of GHR are highlighted. Residues involved in oncogenic mutations leading to constitutive activation [83] are colored red. Protein molecules are shown as semi-transparent surfaces and cartoon representations. (**E**) Sequence alignments of receptor TMDs including juxtamembrane regions. Asterisks indicate receptors with residue numbers of mature proteins lacking signal peptides. TM α-helical residues are underlined, and those at the dimerization interface are colored red, The WSXWS and related motifs are colored green. Residues in the “switch” and Box1 motifs are colored blue. Arrows indicate key residues in the TMDs of cytokine receptors.

For one of these receptors, hPRLR, we generated models of complexes with three different human hormones known to interact with this receptor *in vivo*: prolactin (PRL), somatotropin (GH1), and placental lactogen (CSH1). These models demonstrate many similarities, but also some differences. For example, in all three models, zinc-binding centers are located in the ligand binding site 1. Zn^2+^ ions may link the α1 and α4 helices of hormones (residues H27 and D183 of hPRL, residues H18 and E174 of hGH1 and hCSH1) with hPRLR (residues D187 and H188), in agreement with experimental studies [6]. The ECD-ligand complexes of hPRLR with hGH1 and hCSH1 are rather similar (with RMSD of 0.9 Å) but differ from the complex with hPRL (RMSD of 2.3 Å and 2 Å, respectively). These differences are likely caused by the width differences of hGH1 and hCSH1compared to hPRL. The separation of the two ECDs to accommodate larger ligands increases the distances between the N-termini of TM helices and slightly shifts the helix crossing point toward the C-terminal end. There is no change in the distances between helix ends that interact with JAK2.

An important element of the models is the pair of interacting TM α-helices. These α-helices are rather long (27-36 residues or 40 to 54 Å) (**Figures 4A-C**), consistent with NMR studies [35]. The polar C-terminal part of TMD extends from the membrane into the cytosol, where some hydrophobic residues from helix ends along with Box1 residues interact with JAK2 (**Figure 5**). For example, the L^253^xxxI^257^W^258^ “switch” motif in mouse EPOR forms a rigid connection between TMD and ICD, which is critical for the JAK2 activation upon EPO stimulation [36].

**Figure 5.**
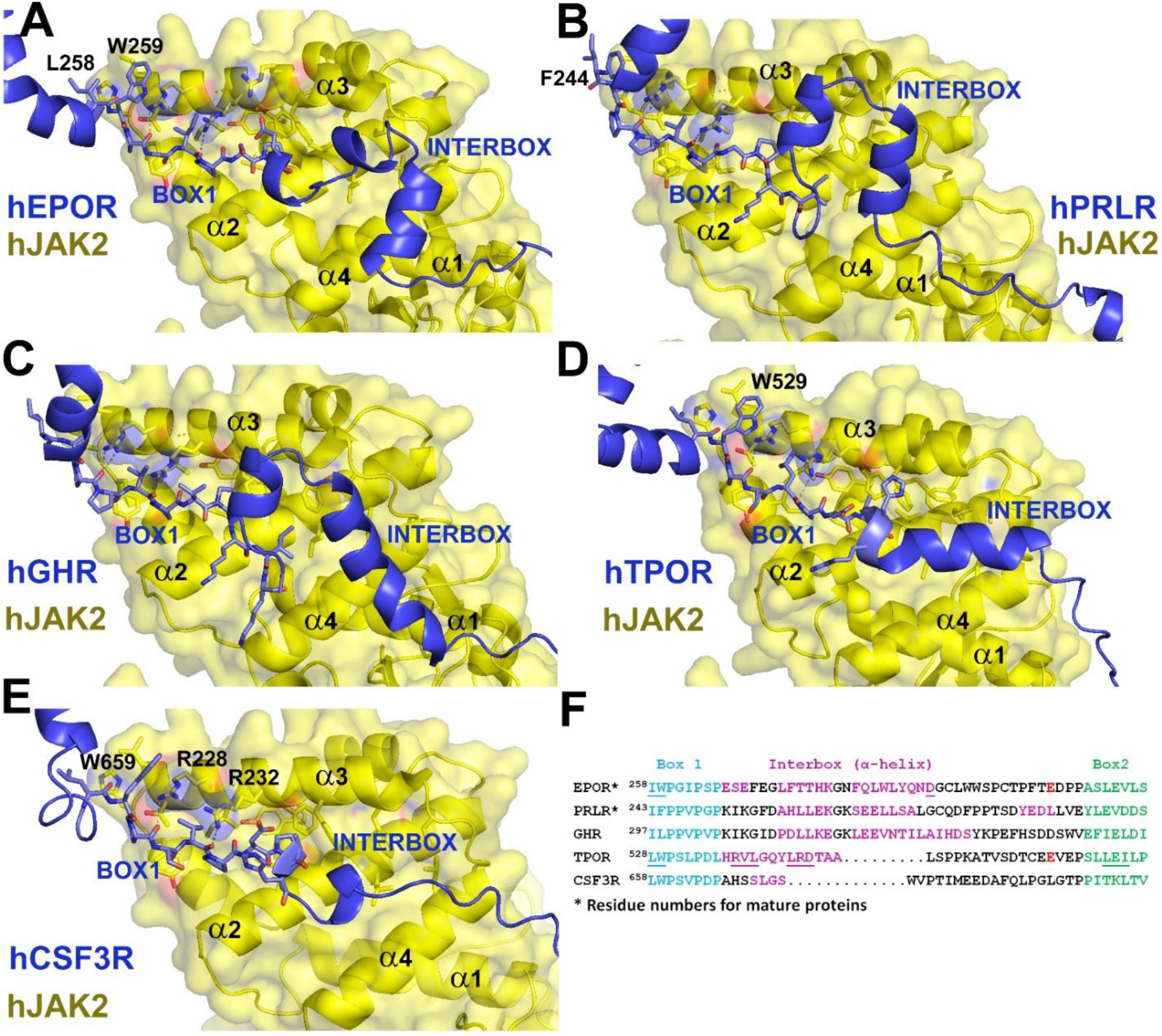
Recognition of receptor Box1 motifs by the FERM domain of JAK2. Fragments of AFM-models generated at step 3 for cytokine receptor TMD-ICD in complex with JAK2. Specific interactions between ICDs of human EPOR (**A**), PRLR (**B**), GHR (**C**), TPOR (**D**), and CSF3R (**E**) and the α1-α4 subdomains of FERM are shown. Pro-rich fragments of the Box1 motifs interact with α3 of FERM; α-helical fragments of the “interbox” regions interact with α2 and α4 of FERM; and Box2 motifs interact with SH2L domain (see **Figure S8**). R228 and R232 from the FERM α3 form H-bonds with the main chain carbonyls of the Box1 fragment. (**F**) Sequence alignments of receptor ICD fragments interacting with the JAK2 FERM-SH2L, based on the AFM models. Residues in the “switch” and Box1 motifs are colored blue. Box2 motifs are colored green, and the interbox regions forming α-helices are colored purple. Underlined residues are known to be important for JAK2 activation [36, 65].

hPRLR is the only homodimeric receptor with a right-handed arrangement of TM α-helices, based on the models of the active receptor dimers (**Table S2**). AFM calculations generated both right-handed and left-handed TM α-helix arrangements for hPRLR complexes with all three ligands with slightly different helix-helix interfaces. However, only the right-handed version appeared in the models of constitutively active hPRLR mutants with deleted ECDs (Δ10-186 and Δ1-210) [37, 38]. The right-handed TMD dimer also had longer TM α-helices than various left-handed dimers. Therefore, the right-handed dimer of hPRLR was selected for further modeling (**Figure 4C**). The selected model is similar to one of conformations of the PRLR dimer obtained in multiscale simulations [39]. The right-handed helix dimer is characterized by a negative crossing angle and the tetrad repeat motif. In the AFM model of hPRLR, TM α-helixes cross at the middle of the membrane at A222, while W214 and W230 of both helices are located near the membrane boundaries. The large distances between N-N and C-C termini of interacting TM helices and the presence of adjacent flexible loops may explain lack of effects of Ala-or Gly-insertions at the junctions of hPRLR TMD with ECD or ICD [40].

In contrast to the right-handed TM dimer of hPRLR, models of hGHR, hEPOR and mEPOR active dimers with bound cytokines demonstrate a left-handed TM helix arrangement, as defined by a positive crossing angle and the (*abcdefg*)*_n_* heptad repeat motif (where the *a* and *d* positions form the interface).

For example, the model of the active hGHR dimer in complex with hGH1 shows a left-handed TM α-helix arrangement with F273 at the *d*-position of the heptad repeat motif (**Figure 4B, Table S2**). This helix orientation and the presence of an intermolecular disulfide C259-C259 are consistent with the NMR structure of the active dimer of TM segments of hGHR [41]. The predicted active conformation of the TM dimer is consistent with Cys-scanning mutagenesis and cross-linking studies that localize residues L269, F273, F276, and V280 at the dimerization interface [42]. The formation of a TM coiled-coil in the active state is consistent with activation of hGHR by the fused coiled-coil dimerization domain of the c-Jun transcription factor that clamps together the TM helices [42, 43].

TM α-helices of the active EPOR dimer also form a leucine zipper with the reference residue, S238 in mEPOR (L239 in hEPOR), occupying the *e*-position of the heptad repeat motif (**Figure 4A, Table S2**). The modeled TM helix arrangement is in good agreement with results of the fusion of the mEPOR TMD with the coiled-coil dimerization domain of the yeast transcription factor *Put3*, where the left-handed dimer *cc-EPOR-III* with S238 in the *e*-position was constitutively active [44]. This helix orientation also explains the constitutive activity of L241N mutation in mEPOR (L242N in hEPOR) [45]. The hTPOR L242N mutated residue is located at the dimerization interface (*d*-position) and may stabilize the TM dimer by forming intermolecular hydrogen bonds.

Despite a generally good agreement of AFM-generated EPOR models with mutagenesis studies, some data cannot be explained by these models. In particular, the AFM EPOR models, as well as corresponding crystal structures are not compatible with the formation of the C129-C129 disulfide that was suggested for the constitutively active R129C mEPOR mutants [46]. In all structures, R129 of mEPOR (R130 of hEPOR) from both receptor chains are located far from each other (Cα-Cα distances of 24-32 Å), and may come closer only if large movements of D2 domains occur in the active state. Such D2 movement is also required to form C132-C132 and C133-C133 disulfides that are suggested for the disulfide-linked constitutively active dimers of E132C and E133C mEPOR mutants [47].

#### TPOR ligand-receptor complex

In the structural model of the human TPO-TPOR (1:2) active complex, TPO binds to D1 (A-B and E-F loops) and D2 (F-G loop) domains *via* multiple hydrophobic and ionic interactions. Five hTPOR residues, F45, L103, F104, D261, and L265, form multiple contacts with hTPO residues from its α1, α3, and α4 helices and the loop between α1-α2 (**Table S4, Figure 6B**). These receptor residues were identified in mutagenesis studies as key TPO-binding determinants [48, 49]. Two receptor ECDs interact not only with the ligand, but also with each other *via* a long loop within D2 (residues 187-238) and two antiparallel β-strands from D4 (residues 436-438) that form a site 3 between D4 domains (**Figure 6A**). Non-conserved cysteine residues from the D2 loop form three intramolecular disulfides (C193-C323, C194-C241, and C211-C322) in many AFM V3 models (**Table S1**). We hypothesize that these disulfides may stabilize a monomeric ECD structure exposed to the extracellular environment, while the D2 loop constrained by disulfide bonds may participate in the ligand binding and in dimer stabilization.

**Figure 6.**
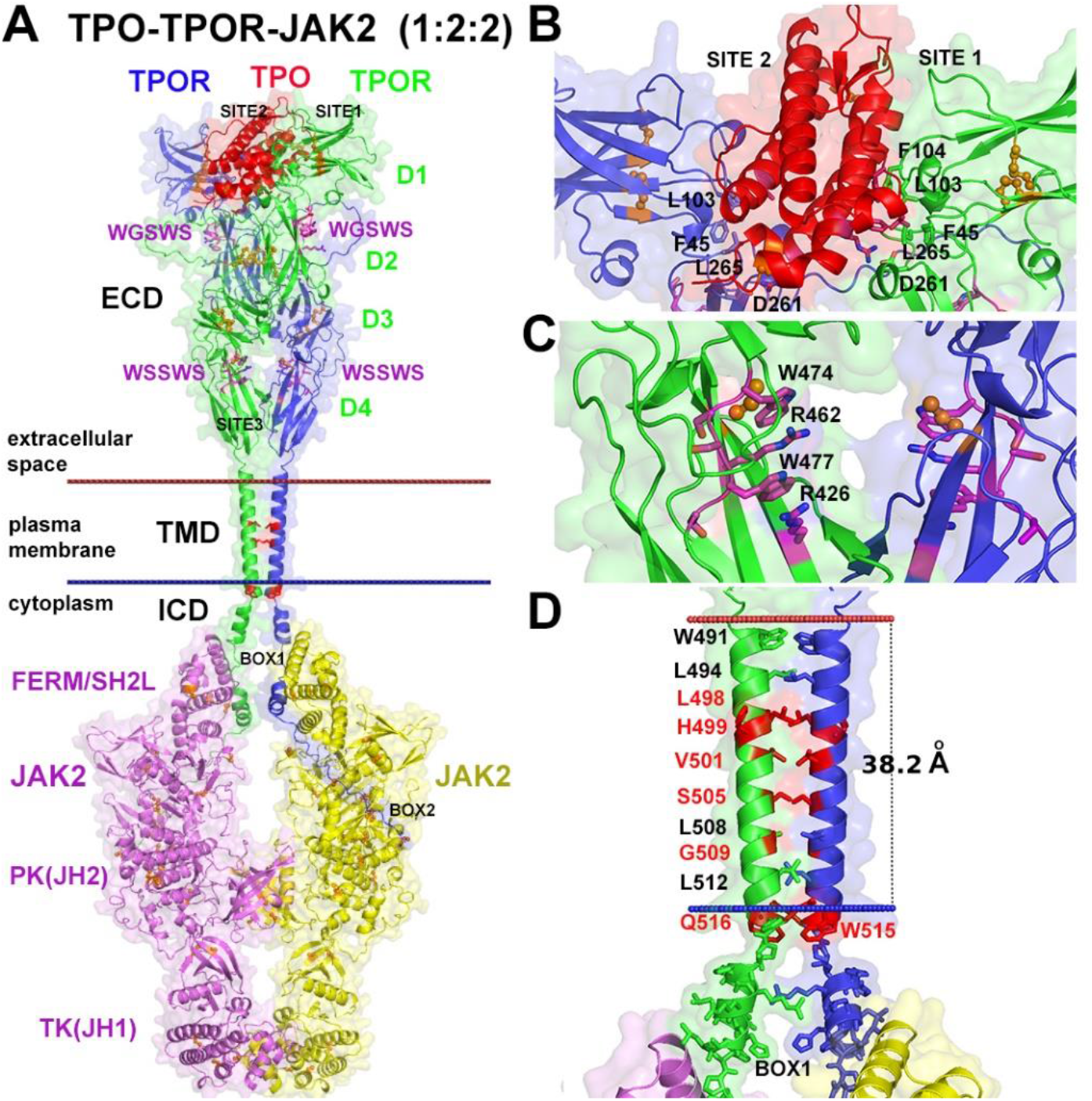
AFM-generated model for the full-length human TPO-TPOR-JAK2 (1:2:2) signaling complex. (**A**) Overview of the complex in a membrane. Protein molecules are shown as semi-transparent surfaces and cartoon representations are colored red for TPO, blue and green for TPOR subunits, and yellow and pink for the JAK2 subunits. Cysteine residues are shown as balls- and-sticks colored orange. Domains of TPOR and JAK2 are indicated. The two WSXWS motifs in D2 and D4 of ECDs, and the Box1 and Box2 regions in ICDs are highlighted. Disease-associated recurrently mutated residues in the TMD (V501, S505, and W515) which cause constitutive TPOR activation [88, 113] are colored red. (**B**) The TPO binding pocket in the ECDs of TPOR model. TPOR residues interacting with TPO are shown as sticks. F45, L103, F104, D261, and L265 have been previously implicated in TPO binding [48, 49]. (**C**) Close-up of the WSSWS motif in the D4 domain of ECD. Aromatic and basic residues involved in the network of cation-π interaction are shown as purple sticks. (**D**) Close-up of the TMD and Box1 region of TPOR interacting with JAK2. TM α-helices of TPOR have left-handed arrangements (with a positive crossing angle); residues at the interface are depicted by sticks, residues with natural or engineered mutations (S505N, L498W/H499C,Y, L498W/W515K, H499G/V501S, H499C,Y/S505N, H499L/G509N, H499L,C,Y/W515K, V501A/W515L,R, S505N/T487A, S505N/S493C, S505N/V501A,M, S505C/W515L, S505N/Q516, and S505N/V501N/A506V) associated with constitutive activation of TPOR [10, 50, 51, 53, 54, 113, 114] are colored red. TPOR residues forming the Box1 motif are shown as sticks.

In the model of the active hTPOR dimer, the TM α-helix spans over 28 residues (from T289 to F517 in the RWQF motif) (**Figure 6C**), similar to long TMDs of human and mouse EPORs [35]. The existence of rather long TMDs encompassing W491 and W515 agrees with NMR studies of TMD dimers [50, 51]. However, a helix break at H499, which has been suggested based on NMR data of hTPOR monomers [52], is not observed in hTPOR, either monomeric or dimeric models generated by AFM. AFM-based models demonstrate the TM α-helix kink at P518. After this helix kink, an additional 3-turn polar helix (A519-L528) extends to the cytosol to interact with JAK2, similarly to the “switch” residues of mEPOR (**Figures 5A, 4D**). The left-handed α-helix arrangement in the model of the active TM dimer with S505 at the *a*-position and H499 at the *b*-position of the heptad repeat motif (**Figure 6D**) is consistent with the dimerization interface of the constitutively active *cc-TPOR-I* fusion construct between a dimeric coiled-coil of Put3 and TMD of mTPOR [53]. This dimerization interface is also supported by Asn-scanning mutagenesis of human and murine TPOR [50] and studies of constitutively active hTPOR mutants [51, 54].

Furthermore, docking of an allosteric ligand eltrombopag to the TM α-helices (**Figure S3**) also supports the proposed AFM model of the active hTPOR dimer. Two eltrombopag molecules are located at the both sides of TM α-helical dimer and participate in hydrophobic interactions with two sets of W491, I492, I494, V495, T496, L498, and H499 residues near the helix N-termini and ionic interactions between drug carboxyl groups and two R456 residues from the D4 domains. Eltombopag can also form Zn^2+^-mediated interactions with both H499 residues, like the structurally similar compound SB394725 [52]. These positions of eltrombopag are consistent with the previously identified locations of its structural analogues [52, 54] and, especially, with a key role for W491 in TPOR activation by eltrombopag [51].

#### CSF3R ligand-receptor complex

Unfortunately, AFM was unable to automatically produce models of ligand-receptor complexes of the long-chain hCSF3R with TMDs forming a dimer, even though the D1-D3 domains with bound ligands were superimposable with the corresponding crystal structure (PDB ID: **2D9Q**)[11] with RMSD of around 3 Å. Therefore, modeling of receptor-ligand complexes for hCS3FR was done stepwise, separately for the ECDs and TMDs. AFM modeling started from a complex of monomeric hCSF3R with bound hCSF3 at 1:1 ratio. Then, two such models were superimposed with both units of the homodimeric cross-over crystal structure of the hCSF3-hCSF3R 2:2 complex composed of two 1:1 units [11]. Superposition demonstrated a good overlap of experimental and calculated 1:1 complexes of D1-D3 domains with hCSF3 (RMSD of 1 Å) (**Table S2**), but different spatial positions of the remaining receptor domains. Small adjustment of the main chain angles in the D3-D4 linker allowed a juxtaposition of TM α-helices to form a dimer.

To define helix orientations in the active ligand-bound state of hCSF3R, we modeled dimers of isolated TM segments with sequences corresponding to native and the constitutively active oncogenic mutant, T640N [55, 56] (**Table S3**). The AFM models of isolated TM segments with native and mutant (T640N) sequences have a left-handed TM helix arrangements with T640 (or N640) at the dimerization interface at the *a*-position of the heptad repeat motif (**Figure 4D**). A similar helix arrangement was predicted by the TMDOCK method [57]. To complete the structure of the full-length active hCSF3R dimer, we combined the model of TM dimers and the model of two multi-domain ECDs with two bound CSF3 ligands. In the final model of the receptor-ligand complex, two gain-of-function mutations, T640N and G644E [55], are located at the TM dimerization interface and can stabilize the TMD dimer *via* hydrogen bonds. The other oncogenic mutations, T612I, T615A, and T118I [55], are located at the D6-D6 dimerization interface (the site 3) and may contribute to stabilization of ECD dimers.

#### Homodimers of ligand-free receptors and TM segments

AFM models of ligand-free receptor dimers were generated for human and murine EPOR and human GHR, PRLR and TPOR. They significantly differ from the corresponding experimental and modeled structures of ligand-bound dimers, with RMSDs ranging from 5 to 9 Å, due to rearrangements of ECDs and TMDS (**Tables S2 and S3**). Unlike the asymmetric active dimers, the ligand-free models are symmetric, have a larger contact area between ECDs of the two receptors chains, a small-sized ligand binding pocket, and altered mutual orientations of TM α-helixes. However, the reliability scores are lower for ligand-free dimer models (ipTM ranging from 0.2 to 03) compared to the ligand-bound dimer models (ipTM ranging from 0.6 to 0.7).

Models of ligand-free homodimers of hPRLR, hGHR, and mEPOR demonstrate a tighter packing of ECDs than in the ligand-bound dimers with occluded ligand binding pockets (**Figure 7**). The rearrangement of ECDs in the ligand-free dimers brings two symmetric D1 domains closer together compared with the active dimer models, and alter the D2-D2 dimerization interface (site 3); thus, some residues from the ligand binding pocket form receptor-receptor interactions occluding the ligand-binding pocket. For example, the close packing of ECDs in the ligand-free hPRLR brings together D187 and H188 residues which form a predicted Zn^2+^ binding site with cytokine ligands, hPRL, hGH1 and hCSH1, in the active structure. A new Zn^2+^-binding site might be formed between two D2 domains of the ligand-free hPRLR (**Figure 7C**). Molecular dynamic simulations of ECDs of hGHR also pointed to the increased contact of subunits of the ligand-free dimer [58]. Extensive contacts between ECDs of two antiparallel receptor chains are also observed in the crystallographic antiparallel dimer of the ligand-free hEPOR (PDB ID: **1ERN**) [59].

**Figure 7.**
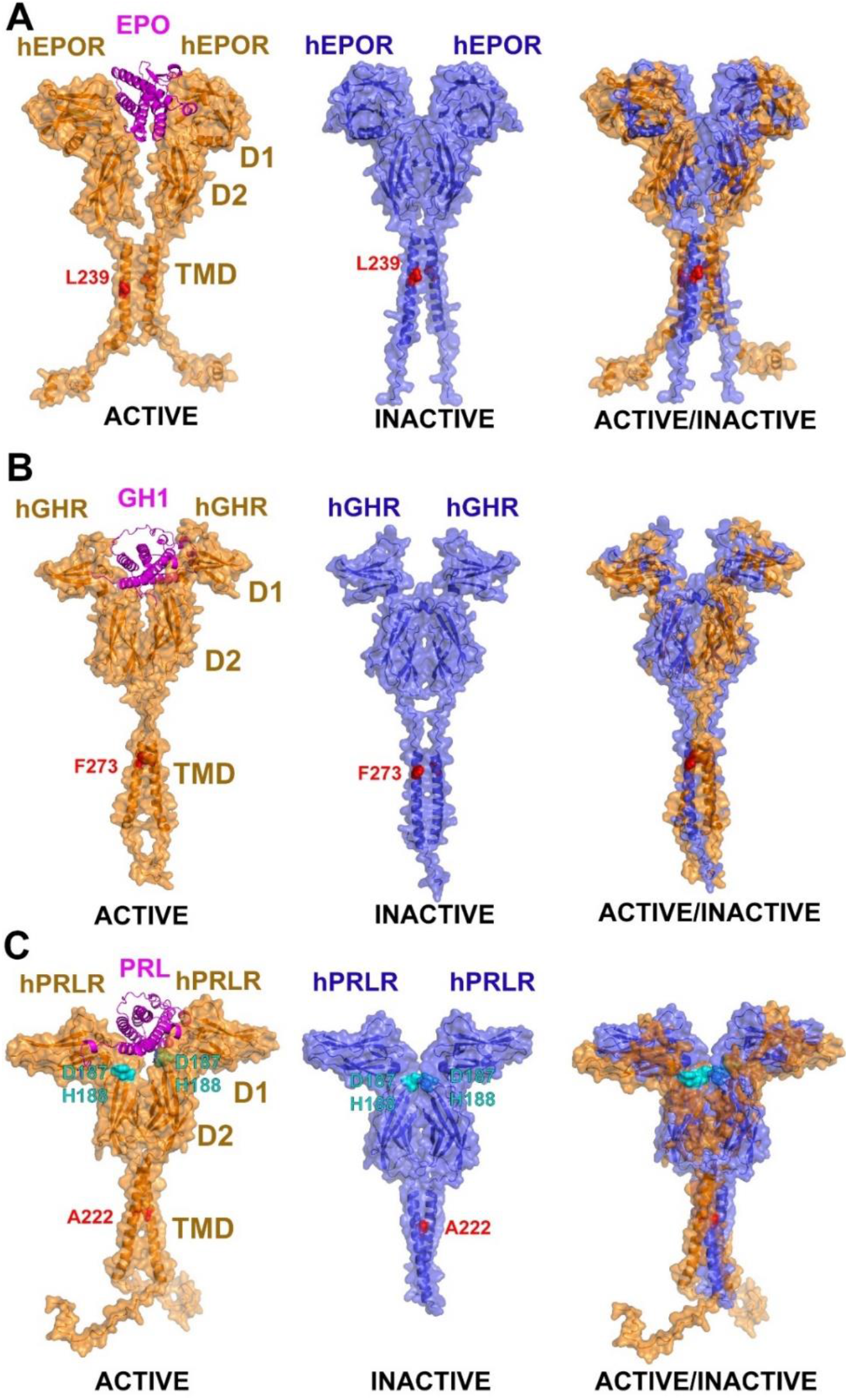
Comparison of AFM-generated models of active and ligand-free dimers of hEPOR (**A**), hGHR (**B**), and hPRLR (**C**). In the ligand-free dimer, D1 domains occlude the ligand binding pockets. Furthermore, the relative positions of D2 domains are changed, and TM α-helix arrangements are different from those in the active dimers. The molecules are shown by cartoon and semi-transparent surface representations colored orange for active dimers and blue for inactive receptor dimers; ligands in the active dimers are colored purple. Reference residues in the TMDs are shown as red spheres. Residue forming a possible Zn^2+^-binding center in hPRLR (D187 and H188) are shown as cyan spheres.

Modeling of the ligand-free hTPOR dimer produced two sets of conformations with an open and closed ligand-binding pocket and a different arrangement of D1-D2 domains in symmetric chains. The closed conformation is too narrow to fit TPO, while the open conformations have a wider space between D1-D2 that can accommodate TPO after slight domain movements to match asymmetric sides of the ligand.

Importantly, for many of the receptors, the two TM α-helices have different mutual orientations in the inactive (ligand-free) and active (ligand-bound) models generated by AFM (**Tables S2, S3, and S5**). For example, in hEPOR and hTPOR models, the TM α-helices are loosely packed and have a right-handed helix arrangement in ligand-free dimers, but form tightly packed left-handed dimers in the ligand-bound states. The model of the ligand-free hPRLR dimer has a left-handed helix arrangement, compared to the right-handed dimer in the ligand-bound state (**Figure 7C**). Less significant difference in TM helix orientations is observed between inactive and active states of EPOR and GHR dimers, where dimerization interface rotates only by ∼50° and ∼100°, respectively (**Table S5, Figures 7A, 7B, S8**).

In addition to the calculations of full-length ligand-bound and ligand-free complexes, we modeled dimers formed by TM-JM peptides (**Tables S3, S5**). Interestingly, the arrangements of TM α-helices in TM-JM peptide dimers are more similar to TM α-helix packing in active (ligand-bound or constitutively active) than in inactive dimers of full-length receptors (**Tables S5**). For example, AFM predicted the same right-handed TM helix arrangements for the full-length active (ligand-bound) hPRLR dimer and its constitutively active mutants, Δ1-186 [37] and Δ1-210 [38], which lack large parts of their extracellular domains.

However, the calculated arrangements of TM α-helices are often close but not exactly the same in dimers of TM-JM peptides and full-length ligand-bound receptors. For example, the model of full-length hTPOR has a left-handed TM α-helix arrangement with S505 at the *a*-position of the heptad repeat motif (**Tables 1, S2**), which has been experimentally proven for the full-length human and mouse TPOR [50] and the left-handed dimer of *Put3*-fused *cc-TPOR-I I* construct [53]. However, S505 occupies an alternative *d*-position the left-handed dimer calculated by AFM for TM-JM peptides of constitutively active hTPOR mutants, L498W/H499Y and H499L/W515K [51], consistent with the helix orientation found in isolated TM helix dimer of the constitutively active S505N mutant [54]. AFM calculations also reproduced two dissimilar TM dimerization interfaces that were identified in *Put3*-fused constructs of mEPOR [44] and its TMD segments [45] with S238 located at the *e-* or *a*-positions of the heptad repeat motif, respectively. Furthermore, AFM predictions of helix orientations in isolated TMDs of hTPOR, mEPOR, and CSF3R agree with low-energy models generated by the TMDOCK method [57] (**Table S5**)

### Step 2: Modeling of the human JAK2 homodimer

An important component of the active signaling complexes are JAK non-receptor kinases that are constitutively associated with ICDs of cytokine receptors. Each member of JAK family is composed of four structural domains: a FERM (four-point-one, ezrin, radixin, moesin) domain, a Src-homology 2-like domain (SH2L), a pseudokinase domain (PK or JH2), and a catalytically active tyrosine kinase domain (TK or JH1) (**Figures 8A, 8B**). JAK2 is the main non-receptor kinase interacting with class 1 homodimeric cytokine receptors. Though experimental crystal structures were obtained for individual domains of JAK2, there is no experimental structure for the full-length JAK2 and its active dimer. Computational models of the full-length JAK2 dimer have been proposed using long-timescale molecular dynamics simulations [60].

**Figure 8.**
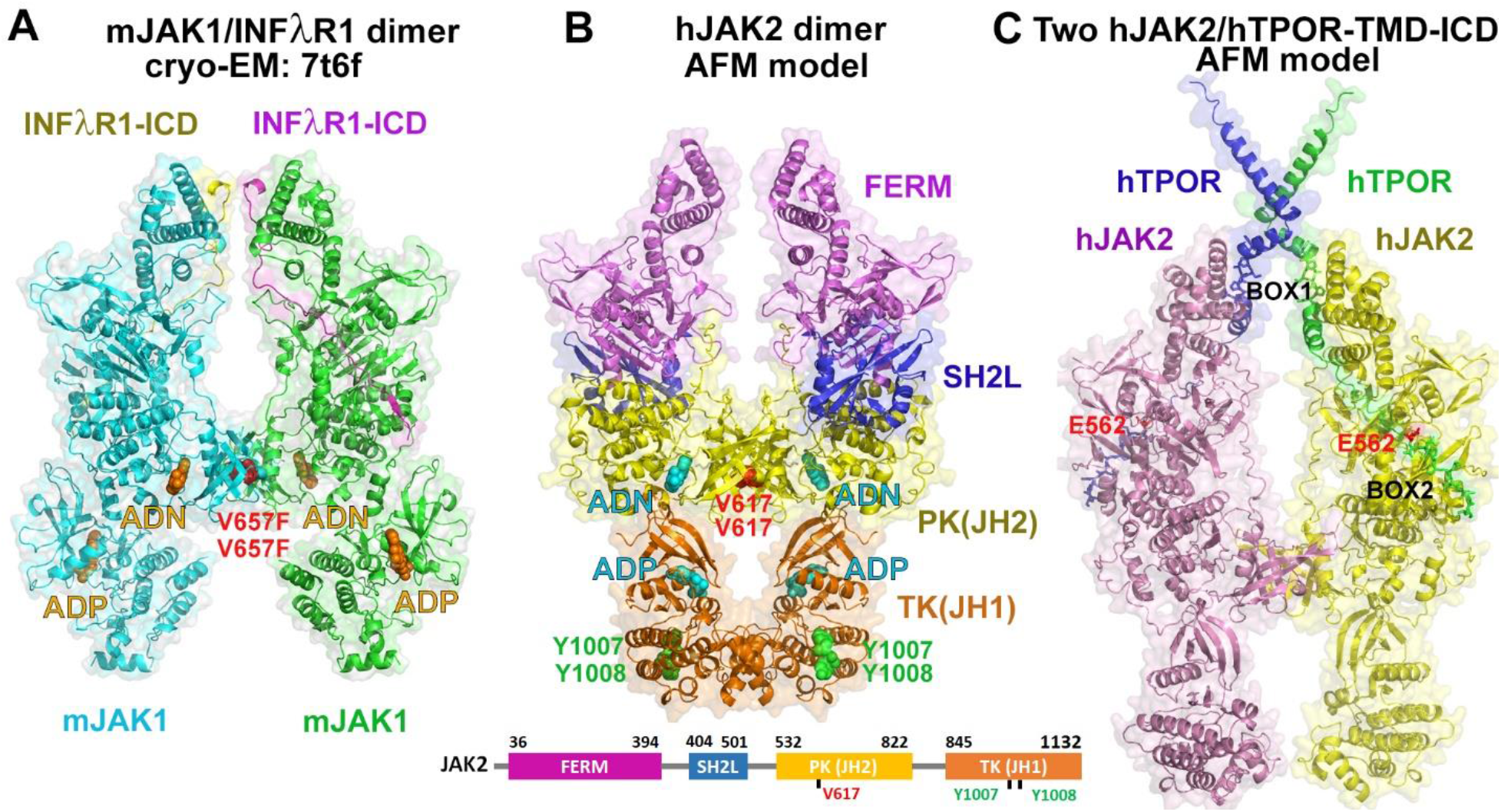
Experimental and computational models of the active dimeric complexes of full-length mouse JAK1 (**A**) and human JAK2 (**B, C**). Two JAK1 or JAK2 subunits dimerize *via* formation of antiparallel β-strands between β-structural N-lobes of PK domains. These domains contain the oncogenic V657F mutation in mJAK1 or the wild-type V617 residue in hJAK2 (shown by red spheres). (**A**) cryo-EM-based model (**PDB ID: 7T6F**) of the mJAK1 dimer in complex with peptides from ICDs of interferon λ receptor1 (INFλR1). JAK1 subunits are colored green and cyan; INFλR1-derived peptides are colored yellow and purple; ligands (adenosine (ADN) and adenosine-5’-diphosphate (ADP)) are shown by orange spheres. (**B**) AFM-generated model of the human JAK2 dimer with colored FERM, SH2L, PK and TK domains. The cyan spheres indicate ADN in PK domains and ADP within TK domains. The green spheres indicate tyrosines (Y1007 and Y1008) from the flexible 15-residue activation loops (residues 993-1017) of TKs that undergo *trans*-phosphorylation during JAK2 activation. The positions of small molecules, ADP and ADN, are similar to those in cryo-EM structure of mJAK1 (A). The lower panel shows a schematic representation of the hJAK2 domain architecture. (**C**) Structure of the JAK2 dimer based on AFM-generated models of two hJAK2 monomers, each in complex with a part of ICD domain of hTPOR. The TMDs of hTPOR are also shown. To form the dimer, the calculated monomeric models were superposed with FERM-SH2L domains of the JAK2 dimer shown in **B**. Each JAK2 subunit (colored yellow and pink) is constitutively bound to the ICDs of TPOR (colored green or blue) *via* Box1 and Box2 motifs. E582 (colored red) of TPOR occupies the aberrant binding pocket for phosphorylated tyrosine in the SH2L domain of JAK2. Protein molecules are shown by semi-transparent surface and cartoon representations.

Recently, cryo-EM structures were obtained for the full-length mouse JAK1 active dimers in complex with the ICD fragments of interferon λ receptor1 (INFλR1) stabilized by the oncogenic V657F mutation (analogous to V617F mutation of JAK2) and nanobodies [61, 62] (**Figure 8A**). Both structures (PDB IDs: **7T6F, 8EWY**) demonstrate that the dimerization interface is formed between β-structural N-lobes of PK domains. The V657F oncogenic mutation stabilizes the dimeric state by participating in a cluster of contacting aromatic residues at the dimerization interface. Interestingly, these dimer structures demonstrate the different relative positions of TK domains connected by the long flexible loops to PK domains that can be closer together or further apart from each other. Such positional flexibility of the TK domains may be essential to facilitate their *trans*-phosphorylation at tyrosine residues from the activation loop, the key step in JAK activation, and for the subsequent phosphorylation of tyrosine residues of associated receptors and STAT proteins.

At the second step of the AFM modeling, dimers of the full-length human JAK2 were generated with and without short ICD fragments of receptors (**Figure 2**, **Table S2**). The presence of the short receptor fragments did not affect the results of the calculations. The models of the hJAK2 dimer were similar to the cryo-EM-based structures of the mJAK1 dimer (**Figure 8B, Table S2**), but the distances between two symmetric FERM domains in the models (L224 Cα-Cα distances), varied from 30 to 60 Å (**Figure 2**). We selected the model of hJAK2 dimer with the minimal distance, similar to 30 Å observed in the experimental structure of mJAK1 dimer (PDB ID: **7T6F**). The dimerization interface in the model was formed by two PK domains, similar to that in the mJAK1 dimer. The dimer is stabilized through association of β-strands connecting the SH2L and PK domains (residues 534-538), two N-lobes and a C-helix of the PK domain. The oncogenic V617F mutation is located at the PK dimerization interface where two V617F residues of the mutant form a cluster with four aromatic residues (F537 and F595 from each JAK2 subunit) which strengthen PK-PK interactions in JAK2 dimer (**Figure S4**).

### Step 3: Modeling of monomeric JAK2 complexes with receptor TMD and ICDs

The third step included building of a complex for each of five receptors composed of a single molecule of JAK2 and receptor TMD and ICD domains; in some cases, a membrane-proximal ECD domain was also included (**Figure 2**). The AFM models of JAK2 monomers superimpose well with the FERM-SH2L crystal structure of hJAK2 [63] and with FERM-SH2L-PK domains of the cryo-EM structure of the mJAK1 dimer [61] or the modeled hJAK2 dimer: the RMSD values were less than 2.5 Å (**Table S2**). Interestingly, a few models generated by AFM-V3 represented the more compact autoinhibited (inactive) conformation of JAK2 with the kinase (TK) domain located close to FERM-SH2L domains and interacting with the PK domain near the kinase active site. This JAK2 domain arrangement is similar to that observed in the crystal structure of PK-TK module of TYK2 (PDB ID: **4OLI**)[64].

It has been assumed that the JAK2 FERM-SH2L domains determines the specificity of receptor binding by engaging the receptor Box1 and Box2 cytoplasmic regions [63]. Indeed, AFM-generated models demonstrate that each receptor interacts with JAK2 *via* the hydrophobic “switch” residues at the TM helix ends, such as L^253^, I^257^, and W^258^ in mEPOR [36], Box1 residues positioned along the α3 of the FERM domain, some α-helical fragments from the interbox region interacting with FERM α2 and α4, and Box2 residues located in the grove in the SH2L domain (**Figures 5, S5**). This membrane-proximal ICD region in cytokine receptors represents the minimal functional core for signal transduction [65]. It was shown that the “PxxPxP” Box 1 motif is essential for binding and activation of JAK2, while the hydrophobic “switch” motif, the acidic and hydrophobic residues from Box2, and several interbox residues are required for JAK2 activation [65] (**Figure 4E**).

Two interbox α-helices from the JAK2-TMD-ICD model of hEPOR (**Figure 5A**) overlapped well (RMSD of 0.9 Å) with the same helices observed in the crystal structure of hEPOR ICD peptide in complex with JAK2 FERM-SH2L domains (PDB ID: **6E2Q**) [66]. Interbox α-helices found in JAK2-TMD-ICD models of hPRLR and hGHR (**Figures 5B and 5C**) are supported by NMR studies of ICD-derived peptides in lipid vesicles [67, 68]. Additionally, in the AFM models for TPOR and EPOR, a glutamic acid preceding Box2 (E562 in TPOR and E301 in EPOR) bind to the aberrant phosphotyrosine binding pocket of SH2L (**Figures 8C, S6**), similar to interactions observed in the crystal structure (PDB ID: **6E2Q**) [66].

The unfolded part of receptor after Box2 is not bound to JAK2 and remains highly structurally flexible, which allows tyrosine residues located in this region to enter in the catalytic site of TK to be phosphorylated (**Figures S5, S7**). Two TPOR tyrosines, Y631 and Y625, were identified as primary and secondary phosphorylation sites, respectively, while phosphorylation of Y591 was shown to participate in receptor downregulation [10]. Interestingly, in the AFM model, the unphosphorylated activation loop of TK (residues 997-1018) partially occludes the catalytic site of the TK domain (**Figure S7**). Therefore, we suggest that activation of the TK domain after its *trans*-phosphorylation could be induced by the movement of the phosphorylated activation loop away from the catalytic site due to electrostatic interactions between phosphotyrosines (pY1007 and pY1008) and adjacent charged residues. Similarly, phosphorylation of the activation loop in receptor tyrosine kinases relieves the inhibition caused by insertion of unphosphorylated loop into the kinase active site [69].

### Step 4: Assembly of the ligand-receptor-JAK2 complexes for five receptors

Assembly of the final structure of the full-length cytokine-receptor-JAK2 complexes included several sub-steps (shown for STEP4 in **Figure 2)**. We first produced the model of the ICD-kinase dimer by superposing two JAK2-TMD-ICD receptor units obtained at the step 3 with FERM-SH2L domains of the JAK2 dimer selected at the step 2. Second, we joined the models of the ICD-kinase dimer and the extracellular ligand-bound complex (selected at the step 1) by superimposing C-termini of their TM α-helices and adjusting conformations of connecting residues between the TM dimer and Box1 motifs. Third, to improve the PK-PK dimerization interface, we replaced PK and TK domains in the final model by the corresponding PK-TK dimeric structure taken from the active JAK2 dimer model (selected at the step 2). Finally, we refined the structures using local energy minimizations with CHARMM c47b2.

The final models of cytokine-receptor-JAK2 complexes are close to the corresponding experimental structures of extracellular receptor complexes and JAK2 dimers (rmsd of common C^α^ atoms was 1.3 to 3 A, **Table 1)**. The models are also consistent with key residues involved in packing of TM α-helices and extracellular domains (**Tables 1, S2-S5**) and other published experimental data, as described in Results (see steps 1 to 3 above) and Discussion.

**Table 1.**
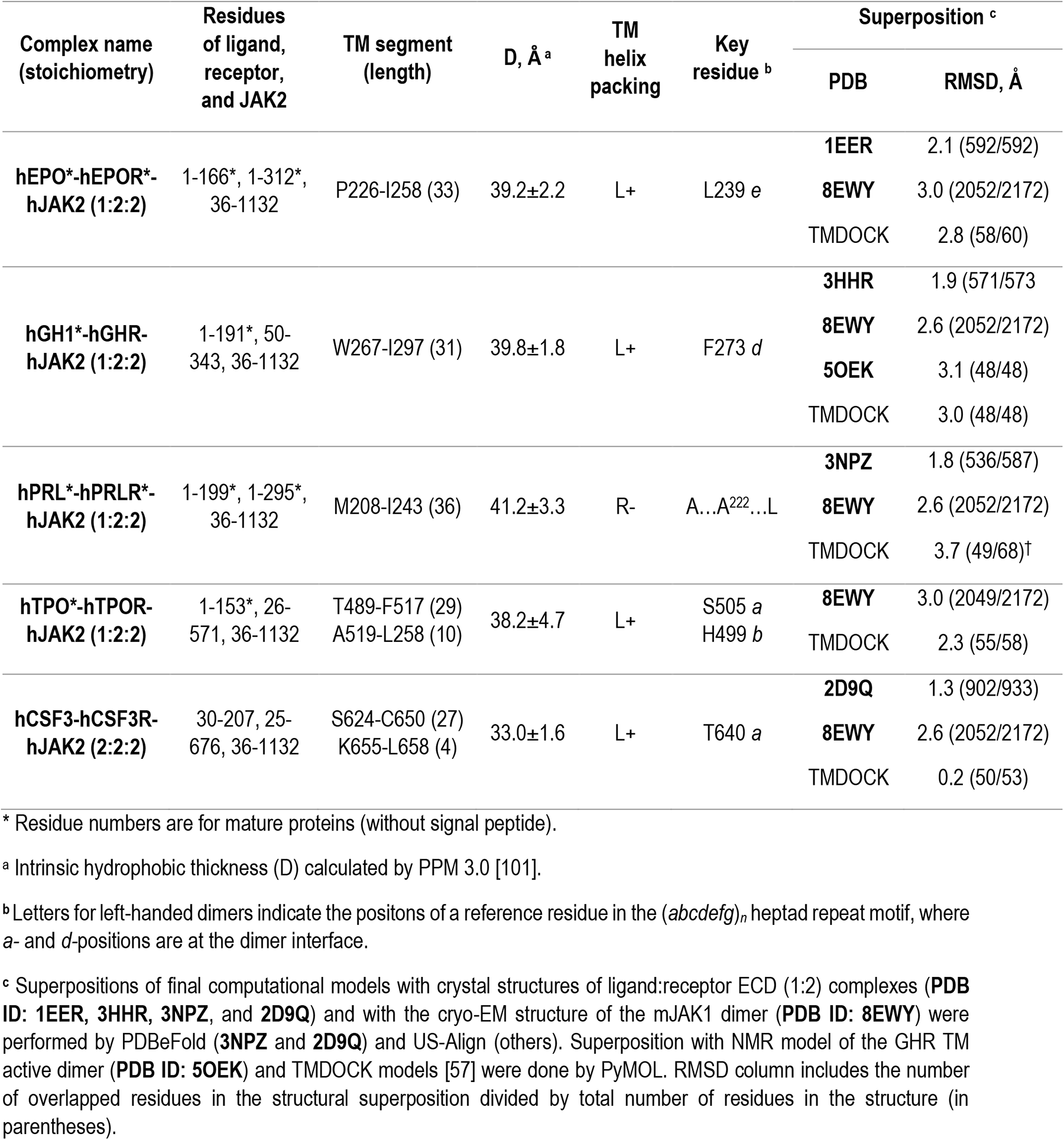
Characteristics of the final models of five homodimeric class 1 cytokine receptor signaling complexes.

### Setting up all-atom MD Simulations

AFM modeling uncovered the conformational heterogeneity of cytokine receptors, especially in the region with low reliability, such as loops connecting protein domains, ICDs, and TM α-helices (**Figure S2**). Though we selected one final model of signaling complexes for each receptor studied (**Figure 1**), other AFM-generated models for active ligand-receptor complexes, as well as inhibited and active JAK2 conformations with different positions of TK domains, activation loops, and JAK2-bound receptor ICDs (**Figures S5 and S7**), may represent different states or snapshots of the conformational dynamics of receptor complexes during their activation. The all-atom molecular dynamics (MD) simulations of these complexes in realistic membranes may shed light on structural transitions between different activation states. Particularly important are the most flexible and the least reliably modeled parts of these complexes, such as loops, ICDs, and ends of TMD regions that may change their conformations upon specific binding of small molecules (e.g. eltrombopag) or interactions with physiologically active lipids (e.g. phosphoinositides) [67, 68].

To demonstrate that our models of five cytokine receptor signaling complexes are suitable for all-atom MD simulations in realistic lipid membranes, we built protein-lipid systems for these complexes in explicit lipid mixture corresponding to the asymmetric mammalian plasma membrane (**Table S6**). In this study, we used the CHARMM force field for proteins and lipids and TIP3P water model [70, 71] with Na^+^ and Cl^-^ ions (see Methods). After successful equilibration of each model in a multicomponent lipid bilayer system, we performed a short production run of 10 ns for each system and deposited the obtained structures together with simulation systems in CHARMM-GUI Archive (https://www.charmm-gui.org/docs/archive/bitopictm). Further studies of structural dynamics of these complexes in the plasma membrane using all-atom MD simulations are beyond the scope of this work.

## Discussion

### AFM-generated structures of signaling complexes of homodimeric cytokine receptors

We have exploited the power of the Alpha Fold Multimer method [32] to generate three-dimensional (3D) models of active ligand-receptor-kinase signaling complexes of five homodimeric class 1 cytokine receptors. The models are consistent, at the level of atomic details, with various experimental data, including available crystal structures of ligand-receptor complexes and cryo-EM-based structures of homologous JAK1 dimers (see Results).

The obtained structural models reveal the quaternary structure of full-length protein complexes. The complexes are well-defined continuous structures extending from ligand-stabilized ECDs of receptors to the large intracellular JAK2 dimer via a long membrane-spanning TMD. Ligand, the cornerstone of a complex, holds together two receptor-kinase units, providing rigidity and stability to the whole structure (**Figure 1**). The first 50-60 residues in the intracellular loops of receptors are bound to a groove on the JAK2 surface and therefore have a fixed structure, while the remaining residues are apparently disordered and flexible, which facilitates the phosphorylation of specific tyrosine residues in the receptors followed by their binding to SH2 domain-containing proteins from the JAK-STAT signal transduction pathway.

Despite the overall similarity of these five signaling complexes, they are different in a number of aspects (**Figure 1**). They have different domain compositions. The receptor-ligand binding stoichiometry is 1:2 for all receptors, but 2:2 for the CSF3-CSF3R complex. Finally, the mutual arrangements of two TM α-helices in the TM dimers are receptor-specific. The significant piston movement of one TM helix relative to the other caused by ligand asymmetry is present only in short-chain receptors where the ligand-binding domains are located close to the membrane. We suggest that this piston movement does not have a significant functional role.

The packing modes of ECDs and TMDs in dimers are receptor-specific and depend on residue compositions, functional states, and upon the nature and stoichiometry of the bound ligands (**Table 1**). For example, the model of the ligand-bound hPRLR complex has a right-handed TM α-helix arrangement, while the packing of TM α-helices in models of the active state of all other receptors is left-handed. Modeling of the same hPRLR receptor with three different ligands demonstrates slightly different helix orientations of the right-handed TMD dimers in the complexes (**Table S2**). Altered TM α-helix arrangements in the active receptor dimer may induce different physiological responses, as shown using various *Put3*-fusion constructs of mEPOR [44] and mTPOR [53].

Furthermore, the mutual arrangements of two TM α-helices are different in models of ligand-bound (active) and ligand-free (inactive) complexes (**Tables S2, S3, S5**).These results are consistent with experimental observations of distinct rotational positions of the TM helices in the active *vs* inactive dimers of mEPOR, mTPOR, and hGHR [42, 44, 53].

### Activation mechanism

The exact molecular mechanism of cytokine receptor activation that triggers JAK2 activation (**Figure 9**) remains a matter of debate. It is accepted that dimerization is essential but not sufficient for receptor activation [72], and that receptor dimerization is driven by the association of the TM α-helices [41, 73–75]. Moreover, receptor activation requires a specific orientation of receptor TM helices to form a productive dimeric state that brings the ICD-bound JAK2 molecules into positions competent to initiate intracellular signaling [44, 53]. However, it remains controversial whether the activation mechanism involves the ligand-induced receptor dimerization (activation model 1, I-III-IV pathway in **Figure 9**) or conformational changes in preformed inactive receptor dimers upon ligand binding (activation model 2, I-II-IV pathway in **Figure 9**) [76].

**Figure 9.**
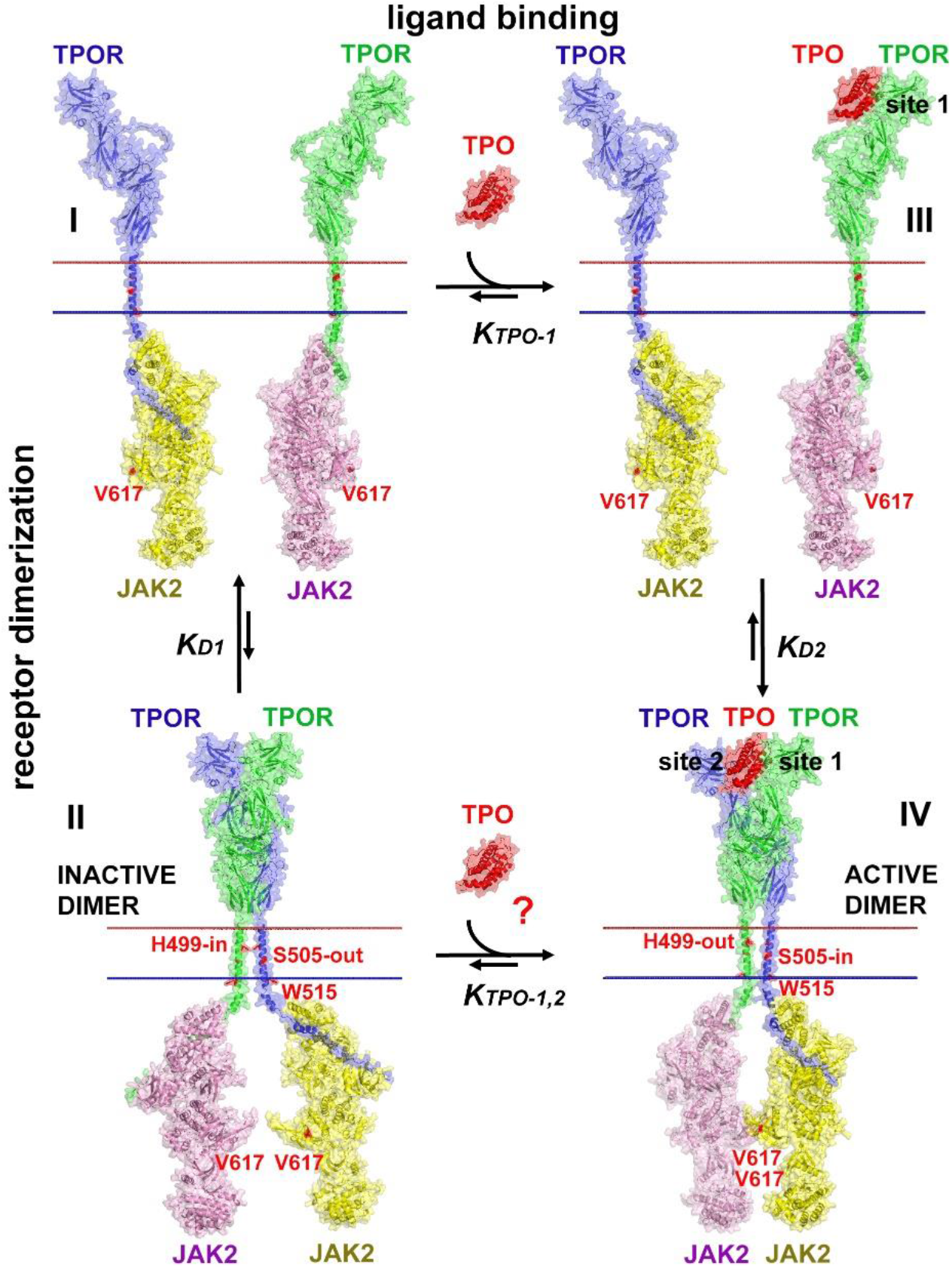
Suggested activation mechanism of homodimeric class 1 cytokine receptors (exemplified by TPOR) based on the AFM modeling and published live-cell dimerization assay [19]. In the absence of the TPO ligand, TPOR receptors are mainly in the monomeric state (state I) [19], even though some dimerization may occur (state II). Ligand binds first *via* site1 to one receptor chain (state III), and then *via* site 2 to the second receptor chain. This leads to stabilization of the active receptor dimer (state IV) with specific rotational orientations of TM α-helices whose intracellular ends bring close two JAK2 molecules bound to receptor ICDs (Box1 and Box2 motifs). This enables the dimerization and activation of JAK2. Protein molecules are shown in semi-transparent surface and cartoon representations, colored red for TPO, blue and green for TPOR subunits, yellow and pink for JAK2 subunits. Residues that are involved in the constitutive activation of receptor (S505, W515) and JAK2 (V617) or those that regulate the formation of the active TMD dimer (H499) are colored red [50, 51, 89, 114, 115].

In the activation model 1, receptor dimerization occurs only in the presence of appropriate ligands. This model has been recognized for many years and gained an additional support in recent studies of dimerization of cytokine receptors in living cells using single-molecule fluorescence microscopy [19]. These studies demonstrated that human TPOR, GHR, and EPOR exist as monomers at the physiological receptor densities in plasma membrane, while the basal dimerization level is negligible. Evaluation of energy contributions showed that binding of TPO to TPOR provides the main contribution to the total dimerization energy. It was also estimated that the intrinsic dimerization affinity of TPOR-JAK2 subunits is low, but constitutively active oncogenic mutations in the dimerization interface of JAK2 (V617F, M335I, H538L, K539L, H587N, C618R, and N622I) and in TMD of TPOR (W515K) provide additive stabilizing free energy contributions which promote TPOR-JAK2 dimerization and formation of the active signaling complex.

An alternative activation model 2 suggests that receptor pre-dimerization occurs in the absence of ligands, and dimer reorganization follows after ligand binding. This model has been proposed based on the extensive structural, biochemical, and mutagenesis studies of different cytokine receptors, including human and mouse EPOR [44, 75], human GHR [42, 72, 77], human PRLR [78], human and mouse TPOR [50, 51, 53], and their TMD fragments [41, 45, 54]. Binding of a specific ligand to the inactive preformed dimer is required for conformational changes and reorientation of receptor TM α-helices to form an active dimeric state that induces proximity and dimerization of associated JAK2 subunits [76].

There are several experimental observations that challenge the hypothesis of ligand binding to pre-existing dimers (model 2). First, at physiological concentrations of receptors at cell surface, the fraction of monomeric receptors (state I) is much higher than of preformed dimers (state II) [19]. Second, the bell-shaped dose-dimerization curve [10, 19] is consistent with the two-step ligand binding to monomeric receptors (state III): initially *via* site 1 to one chain, then *via* site 2 to the recruited second chain, which leads to formation of the ligand-receptor complex (state IV). This dimerization is inhibited by the presence of excess ligand that binds *via* the high-affinity site 1 to receptors, blocking further receptor dimerization via site 2 interactions.

AFM-based modeling provides insights into possible activation mechanisms. The modeling uncovered that ligand-free dimers could be formed for many receptors, but such structures have occluded ligand binding pockets incapable of accommodating large cytokine molecules, along with an unproductive arrangement of TM helices. This is in line with the notion that ligand-free ECDs lock receptors in the inactive states, as PRLR and TPOR variants lacking large parts of ECDs are constitutively active [78, 79]. Ligand binding to preformed dimers with the closed ligand binding pocket would require a significant rearrangement of their ECDs and TMDs and possibly even the dissociation of two receptor molecules. Based on these findings, AFM modeling generally provides more support for model 1 of the ligand-induced dimerization and activation, the I-III-IV pathway (**Figure 9**), at least for class 1 homodimeric cytokine receptor complexes.

Nonetheless, ligand binding to preformed dimers (I-II-IV pathway, **Figure 9)** could represent an alternative non-canonical activation pathway that is used by some receptors in particular cases. For example, we have recently proposed the two-step TPOR activation by the MPN-associated calreticulin mutants (CRTmut) [20] that are suggested to form a CRTmut-TPOR (2:2) complex [80]. The formation of the CRTmut-TPOR active complex takes place in endoplasmic reticulum (ER) membranes where the local density of preformed dimers of immature TPOR molecules may be relatively high. We propose that at the first step, a dimer of CRT mutants binds to the preformed TPOR dimer with the occluded ligand binding pocket *via* interactions with immature mannose-rich glycans linked to N117 residues of both receptor chains. Then, as the second step, CRTmut ligands induce rearrangement of receptor ECDs followed by the insertion of positively charged C-terminal helices of CRTmut into the unlocked binding pocket between D1-D2 domains of both TPOR chains. A generally similar model for the final active CRTmut-TPOR (2:2) complex has been proposed based on a combination of experimental and computational approaches [21]. The CRTmut-TPOR (2:2) active complex formed by ligand binding to the preformed receptor dimer, together with ICD-associated JAK2 traffics from ER to the plasma membrane *via* the secretory pathway [81]. Whether other cytokine:receptor complexes might also use the I-II-IV pathway (**Figure 9)** remains to be further examined.

Importantly, even the ligand-induced dimerization pathway (I-III-IV) implies significant structural changes during the formation of an active signaling complex. The rotational and translational movements of both ECDs relative to each other and side chain rotations are required to adjust the binding pocket for an asymmetric ligand. However, such molecular movements do not change the overall structure of individual monomeric subunits because they are nearly identical in the active and inactive dimers (RMSD <0.6 Å). This rotational and piston motions propagate toward the membrane leading to a receptor-specific arrangement of TM α-helices and adjacent ICD Box 1 residues to bring together ICD-bound JAK2 molecules in an orientation appropriate for productive JAK2 dimerization, activation, and subsequent triggering of signaling events. To the contrary, in the ligand-free inactive dimers, receptor-associated JAK2 subunits are spatially separated and have incorrect orientations, which prevents their dimerization.

The results of the modeling are consistent with FRET studies of hGHR signaling [42]. In the ligand-free inactive conformation of the GHR-JAK2 (2:2) complex, JAK2 subunits allow FRET reporters (mCit and mCFP) covalently attached to the receptor C-termini (37 residues below the Box1 motif) to approach each other at the distance of ∼47 Å from the one side of the JAK2 dimer (**Figure S8A**). The active GH1-GHR-JAK2 (1:2:2) complex have the TM dimer interface different from that in the inactive dimer by ∼100° rotation of F273 towards the dimerization interface (from *e-* to *d*-position of the heptad repeat motif). This TM helix rotation promotes JAK2 dimerization *via* PK-PK interactions. The tightly packed JAK2 dimer prevents FRET reporters coming closer to each other (the distance between chromophores is 75 Å) (**Figure S8B**). Separation of FRET reporters in the active GHR-JAK2 signaling complex is in agreement with experimental data [42].

### Mapping of oncogenic mutant onto signaling complexes

Mutations of cytokine receptors and JAK2 have been implicated in dysregulation or chronic activation of cytokine pathways leading to severe pathologies, including hematological malignancies, growth abnormalities, and aberrant immune responses [5, 81–85]. Disease-associated mutations can be classified as loss-of-function (LOF) and gain-of-function (GOF) mutations. The latter usually cause the constitutive activation of cytokine receptors and JAK kinases [10, 83, 86]. Mapping of known oncogenic missense mutations onto the AFM-based models of active ligand-receptor-kinase complexes may shed light into possible molecular mechanisms of pathological effects of these mutations.

The majority of GOF mutations in JAK2 [86] are located in the PK domain, regions involved in the PK-TK inhibitory interface (**Figure 10**, spheres colored cyan) and PK-PK dimerization interface (**Figure 10**, spheres colored blue). Mutations in the PK-PK dimerization interface are mainly associated with MPNs, with a single point mutation, V617F, identified in more than 95% of polycythemia vera (PV) cases and 50-60% of essential thrombocythemia (ET) and primary myelofibrosis (PMF) cases [87]. V617F together with adjacent aromatic residues, F537 and F593, form a hydrophobic cluster of six aromatic residues from both subunits that stabilizes the JAK2 dimer (**Figure S4**). Other oncogenic mutations are found near this cluster (M535I, H538L, K539L, and N622I). They strengthen the hydrophobic interactions at the dimerization interface. Indeed, all these mutations induce ligand-independent activation and dimerization of the receptor forming the signaling complex with JAK2 [19]. Mutations of the PK-TK inhibitory interface are located between N-lobes of PK and TK. The interface is formed by hydrophobic and charged residues, including the R683-D873 pair. Mutations of interfacial residues, including this ionic pair, can weaken PK-TK interactions and facilitate movement of TK from the inactive (**Figure 10A**) to the active conformation (**Figure 10B**). Activation of JAK2 due to relieved inhibitory function of the PK domain represents a possible mechanism that triggers MPNs, acute myeloid leukemia, and acute megakaryoblastic leukemia caused by these mutations [86].

**Figure 10.**
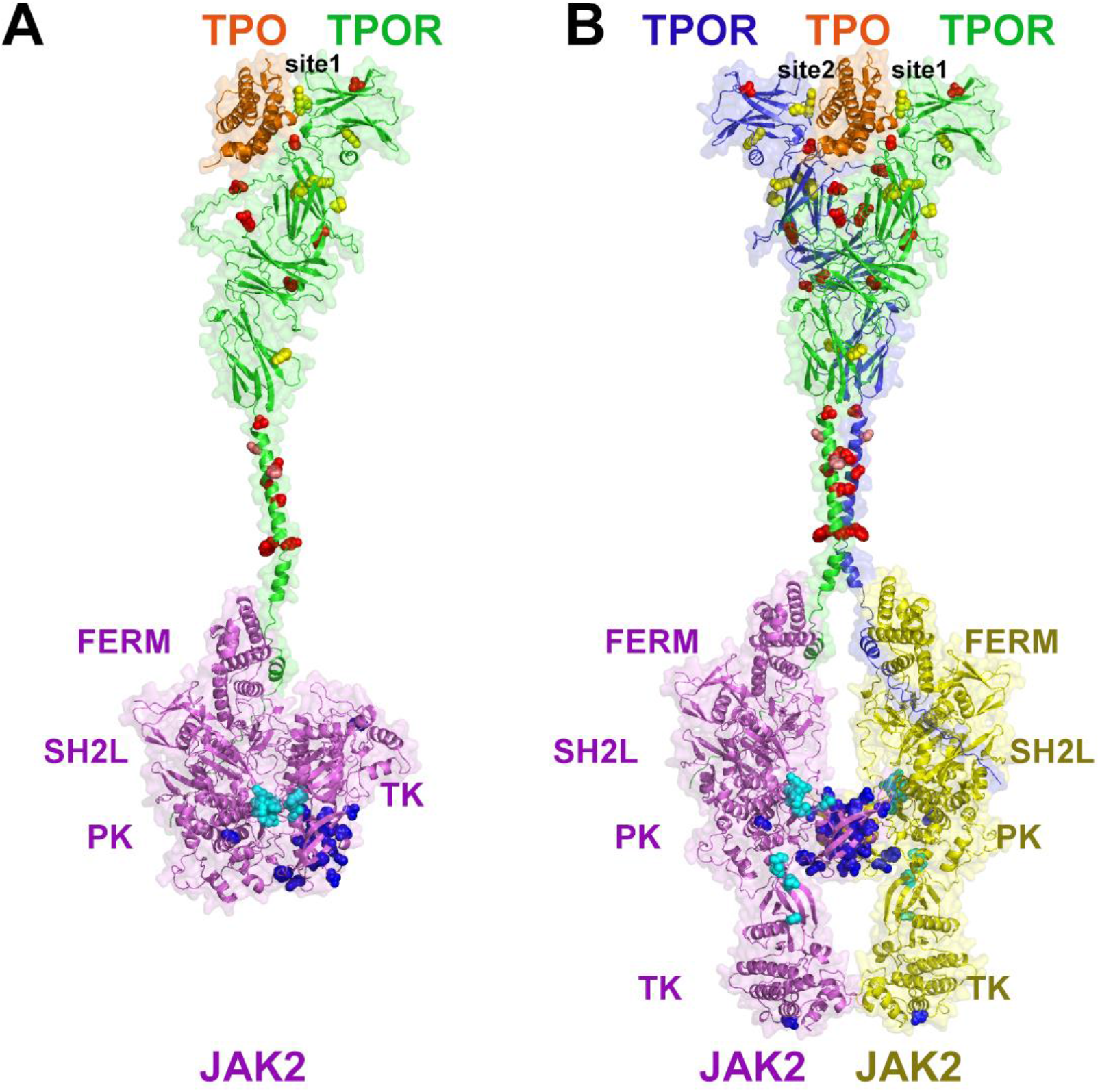
Mapping of disease-associated mutations onto AFM models of TPOR complexes. (**A**) TPO-TPOR-JAK2 complex (1:1:1). TPO binds to TPOR via its high-affinity site 1, forming a binary (1:1) inactive complex. JAK2 is constitutively associated with the ICD of TPOR. JAK2 is in the autoinhibited (inactive) form with the PK domain interacting with the TK domain near the kinase active site, which inhibits the TK’s catalytic activity. (**B**) Active signaling complex of TPO-TPOR-JAK2 (1:2:2) with TK in the active state. TPOR dimer is in the TPO-bound (active) conformation with TMDs forming the left-handed dimer. TPOR mutation sites associated with myeloproliferative neoplasms (MPNs)[10] are shown as yellow spheres for lost-of-function mutations, red spheres for gain-of-function mutations, and pink spheres for ‘enhancer’ mutations in the TMD (S493, H299). MPN-associated mutations of JAK2 are shown as blue spheres for mutations at PK-PK dimerization interface and cyan spheres for mutations at PK-TK inhibitory interface. Molecules are shown as cartoon and semi-transparent surface representations colored orange for TPO, blue and green for TPOR subunits, and yellow and pink for JAK2 subunits.

Oncogenic hTPOR mutations are found in all receptor domains, with LOF mutations located mainly in ECD and ICD and GOF mutations clustered at dimerization interfaces created by TM α-helices and loops inside the ECD (**Figure 10**). LOF mutations usually induce thrombocytopenia, while GOF mutations are mainly associated with thrombocytosis and MPNs, such as PMF and ET [88]. LOF mutations, including K39N, R102P, P106L, W154R, R257C, P635L, show low cell-surface expression, possibly due to defects in receptor folding or trafficking [10]. Another LOF mutation, F104S located in the ligand-binding pocket, impairs TPO binding to the ECDs [10, 88]. The most common GOF mutations identified in MF and ET patients, S505N and W515K/L/A/R, are located within the TMD [10]. These mutations cause constitutive activation of TPOR due to stabilization of the productive TMD dimer. There are also several “enhancer” mutations in TM α-helices that stabilize the active mode of helix dimerization [51, 89], which may enhance the pathological effect.

A more detailed analysis of GOF and LOF mutations in the context of compete structures of active signaling complexes will add to our understanding of the role of disease-associated mutations in cytokine-induced JAK-STAT signaling cascades. Knowing the molecular mechanisms of oncogenic mutations will guide the development of new cancer therapeutic agents.

## Conclusions

Using the transformative ability of the Alpha Fold Multimer to predict structures of proteins and their complexes with high accuracy, we generated models of full-length active signaling complexes for human homodimeric cytokine type 1 receptors, EPOR, GHR, PRLR, TPOR, and CSF3R. Analysis of the resulting models of signaling complexes, as well as models of inactive dimers, examines, in a structural context, highly debated questions related to the mechanism of cytokine-initiated activation that triggers JAK-STAT signaling pathway in cells.

First, we demonstrate that although ligand-free receptors may form stable inactive dimers, the ligand binding pocket in such dimers is occluded, thus preventing ligand binding. At the low cell-surface receptor densities, cytokines are more likely to bind and activate monomeric receptors *via* a two-step process: ligand-induced receptor dimerization accompanied by conformational rearrangements in ECDs and TMDs. This may represent the canonical receptor activation mechanism. A non-canonical activation route through ligand binding to preformed inactive dimers can also occur in specific cases, such as activation of TPOR by the oncogenic CRTmut [20].

Second, we can picture the complete process of JAK2 activation. The process starts from the receptor-induced proximity of two JAK2 molecules, followed by dissociation of PK-TK inhibitory complex in each JAK2 molecule. Subsequently, the dimerization of two symmetric PK domains stabilizes the JAK2 dimer leading to *trans*-phosphorylation of both TK activation loops and their movement away from the TK active sites, enabling tyrosine phosphorylation of receptors and other associated proteins.

Many other aspects of receptor conformational dynamics were also clarified, including atomic details of a specific binding of receptor Box1 and Box2 ICD motifs to JAK2, the ancillary role of the piston TM helix movement in short-chains cytokine receptors, and the role of GOF mutations in stabilizing dimerization interfaces in receptor and JAK2 molecules. The mode of interactions of two molecules of eltrombopag, an FDA-approved TPOR agonist, with the TM α-helical dimer of hTPOR is proposed.

The computational modeling described in this study also uncovers certain limitations of the AFM method. The current versions of the AFM program do not allow a direct modeling of large complexes of multi-domain proteins. Therefore, such complexes need to be assembled from the smaller AFM-generated parts. Moreover, in case of multiple alternative models produced by AFM, the selection of the correct structures still requires supporting experimental data. When such data are lacking or insufficient, modeling of complexes also needs to include comparative analysis of models obtained for sequences of different lengths, with different sets of structural domains, mutants, and subunitstoichiometries. It is anticipated that future versions of the AFM program will overcome some of these limitations, allowing predictions of multiprotein complexes directly and with the improved accuracy.

Despite the limitations, the computational approaches used in this work can be applied in the future to modeling of cytokine receptor complexes from other families, as well as other large functional assemblies of single-pass TM proteins that trigger different intracellular pathways. Knowing 3D structures of such complexes is critical for development of new drugs and therapeutic strategies.

## Methods

### Modeling of signaling complexes with AlphaFold-Multimer (AFM)

Modeling of active signaling complexes of five cytokine receptors was performed using AlphaFold_2.0_multimer.v2 (1.3.0.version) and more recent AlphaFold_2.0_multimer.v3 (1.5.2.version) [32] implemented through ColabFold notebook [33]. ColabFold was downloaded from Github (https://github.com/YoshitakaMo/localcolabfold) together with the environmental databases (https://colabfold.mmseqs.com) and installed on a local computing cluster. ColabFold was running locally using 12, 24, and 48 recycles, MMseq2 for multiple sequence alignments, refinement with Amber, and no templates. The quality of structural models was characterized by the mean of per residue pLDDT score (predicted Local Distance Difference Test) ranging between 0 and 100 that characterizes local structural accuracy [22, 90], as well as using PAE (Predicted Aligned Error) or PAE-derived pTMscore (predicted TM-score) ranging from 0 to1 [91], which correspond to overall topological accuracy. The confidence of the predicted protein-protein interface is assessed by the interface pTM-score (ipTM) ranging from 0 to 1 [26]. For each run, 5 models were generated and ranked by ipTMscores (**Tables S2, S3)**.

The amino-acid sequences from UniProt [92] were used for modeling the following proteins: human JAK2 (UniProt AC: **Q60674**), five human receptors, hEPOR, hTPOR, hGHR, hPRLR, and hCSF3R (UniProt ACs: **P19235, P40238, P10912, P16471**, and **Q99062**, respectively), mouse EPOR (UniProt AC: **P14753**), six human cytokines, hEPO, hTPO, hGH1, hPRL, hCSH1, and hCSF3 (UniProt ACs: **P01588, P40225, P01241, P01236, P0DML2**, and **P09919**, respectively) (**Figure S1**), and mouse EPO (UniProt AC: **P07321**). For hTPO, only the erythropoietin-like N-terminal domain (residues 22-174) [10] was modeled, while the glycan domain was omitted. For an easy comparison with published experimental data, sequences of mature proteins (lacking signal peptides) were used for hEPOR, mEPOR, hPRLR, and cytokines, while full-length sequences of immature proteins (carrying signal peptides) were used in all other cases. In some calculations, especially for bigger complexes, unstructured regions of receptor ICDs below Box1 or Box2 motifs were excluded. For four receptors, except PRLR, one natural ligand was used (**Figure S1**). For human PRLR, which can be activated by three human hormones, prolactin (PRL), somatotropin (GH1), and placental lactogen (CSH1) [6], three receptor-hormone pairs were modeled (**Table S2**). The ligand-receptor stoichiometry was 1:2 for all receptors, except CSF3R, for which the 2:2 complex was modeled.

Additionally, AFM was used to obtain models for ligand-free receptor dimers and receptor fragments composed of TM α-helix with juxtamembrane regions, WSxWS motifs, or adding membrane-proximal domains (D4 for TPOR, D6 and D5-D6 for CSF3R). The complete signaling complexes consist of ligands, receptors, and JAK2 at stoichiometry 1:2:2 (for EPOR, GHR, PRLR, and TPOR) or 2:2:2 (for CSF3R).

AFM modeling was performed in 4 steps (see Results, **Figure 2**). At steps 1-3, more than 30 models were generated for every complex using V2 and V3 versions of AFM with different random seed numbers. Each run with a specified seed produces 5 different models, as illustrated on **Figures S9 and S10** for short-chain dimeric receptors (EPOR, GHR, and PRLR). These models demonstrated similar structures of ECDs but a more variable arrangement of TM helices, and a few of them represented non-interacting monomers. Models with tightly packed TM α-helices were compared with experimental data to select structures most compatible with these data (**Figure 2**, **Table S2**). We found that V3 version produced better models for TPOR complexes (**Figure S11A**), but not for other receptors where the results with V2 and V3 versions were rather similar. Calculations of full-length CSF3:CSF3R (2:2) tetrameric complexes produced models with spatially separated TM helices (**Figure S11B**). Therefore, these complexes were produced by a two-step procedure (see Results).

The final model of each signaling complex was refined using local energy minimization with CHARMM c47b2 [93] through the pyCHARMM python module [94] to remove atom hindrances. The minimization was conducted for 1000 steps (dielectric constant ε=63) with Cα fixation using CONS HARM command in CHARMM (the force of 20), and fixation of cysteines using CONS FIX command. The structures were prepared using the PDB input manipulator on CHARMM GUI [95, 96] and then converted back using MMTSB convpdb.pl [97]. The models of all complexes are available through the Membranome database [98].

AFM-generated models were superimposed with each other and with available experimental structures by PyMOL (www.pymol.org), PDBeFold (SSM) server [99], and the US-align web server [100]. Membrane boundaries were calculated by the PPM web tool [101]. All figures were generated by PyMOL.

### Modeling of lipid bilayer systems with signaling complexes

The final models of cytokine receptor signaling complexes were embedded into the lipid bilayer composed of explicit lipids forming the asymmetric mammalian plasma membrane [102]. The inner membrane leaflet was composed of phosphatidylcholine (PC), phosphatidylethanolamine (PE), phosphatidylinositol (PI), phosphatidylserine (PS), phosphatidic acid (PA), sphingomyelin (SM), and cholesterol (CHOL), while the outer membrane leaflet was composed of PC, PE, SM, CHOL, and glucosylceramide (GlcCer) (**Table S6**). The TIP3P water model was used to simulate explicit water molecules, while the number of ions (Na^+^ and Cl^-^) incorporated corresponded to the physiological concentration (150 mM NaCl). Initial membrane structures were built using the CHARMM-GUI *Membrane Builder* [103–105]. The simulations were performed with all-atom CHARMM36m force field [106], and executed utilizing OpenMM [107]. Following the default equilibration protocol of CHARMM-GUI [93, 96, 108], we first applied NVT dynamics with a time step of 1 femtosecond (fs) for 250 picoseconds (ps). Subsequently, we employed the NPT ensemble with a time step of 1 fs and then with a time step of 2 fs. During the equilibration processes, the protein, lipid, and water molecules were subjected to restraint potentials of their position and dihedral angles. The force constants associated with these potentials were systematically decreased over time. 10 nanoseconds (ns) production runs were performed for each system utilizing a time step of 4 fs and employing the hydrogen mass repartitioning technique [109] in the absence of any restraint potentials. The SHAKE algorithm was employed for managing bonds involving hydrogen atoms [110]. van der Waals interactions were truncated at a cutoff of 12 Å, with a force-switching function applied between 10 and 12 Å [111], while electrostatic interactions were calculated using the particle-mesh Ewald method[112]. The manipulation of temperature and pressure (at a standard pressure of 1 bar) was achieved by utilizing Langevin dynamics with a friction coefficient of 1 ps^−1^ and a semi-isotropic Monte Carlo barostat, respectively.

## Abbreviations

AFM: AlphaFold Multimer
CHM: cytokine homology module
CRT: calreticulin
CSF3: granulocyte colony-stimulating factor
CSF3R: granulocyte colony-stimulating factor 3 receptor
CSH1: placental lactogen or chorionic somatomammotropin hormone 1
ECD: extracellular domain
EPOR: erythropoietin receptor
ET: essential thrombocythemia
FERM: four-point-one, ezrin, radixin, moesin
FnIII: fibronectin type III
GH1: somatotropin
GHR: growth hormone receptor
GOF: gain-of-function (mutation)
JAK2: Janus Kinase 2
ICD: intracellular domain
IL: interleukin
LOF: loss-of-function (mutation)
MD: molecular dynamics
MPN: myeloproliferative neoplasms
NVT: dynamics, constant number, constant-volume, and constant-temperature simulation
NPT: ensemble, isothermal-isobaric ensemble
PK: pseudokinase
PMF: primary myelofibrosis
PRLR: prolactin receptor
PV: polycythemia vera
SH2L: Src-homology 2-like SHP, tyrosine phosphatase
SOCS: suppressors of cytokine signaling
TPOR: thrombopoietin receptor
TK: tyrosine kinase
TM: transmembrane
TMD: transmembrane domain
TM-JM: transmembrane and juxtamembrane regions.

## Data availability

Our models of five cytokine-receptor-JAK2 complexes in explicit lipids of the mammalian plasma membrane and simulation systems are available in CHARMM-GUI Archive (https://www.charmm-gui.org/docs/archive/bitopictm).

## Acknowledgments

This work was funded by the Division of Biological Infrastructure of the National Science Foundation (Award # 1855425 for I.P. and A.L.) and (Award # 2011234 for W.I.) and by the National Institutes of Health (R01AI123957 for M.R.). The authors thank S. Todd for local installation of AphaFold Multimer ColabFold version and design of a web server that was used during this work.

## Supplementary Material

Supplementary Material: 11 Figures demonstrating structures of class 1 cytokines and molecular details of AFM models of cytokine receptor complexes supported by experimental data and 6 Tables presenting quality metrics of different AFM models in comparison, characteristics of receptor partners, interaction residues in the binding pocket of TPOR, and lipid composition of the mammalian plasma membranes (PDF file).

## Notes

https://www.charmm-gui.org/docs/archive/bitopictm

### Competing Interest Statement

The authors have declared no competing interest.

