## Supplementary materials for "Structural modeling of cytokine-receptor-JAK2 signaling complexes using AlphaFold Multimer"

#### Content

**Page 2** **Figure S1**. Structures of class 1 cytokines

**Page 3**. **Figure S2.** Five AFM-predicted models colored by per residue pLDDT score

**Page 4.** **Figure S3**. Suggested binding mode of eltrombopag to TPOT TMD

**Page 5.** **Figure S4.** Close-up of the PK dimerization interface of JAK2 active dimer

**Page 6.** **Figure S5**. Interactions between TK domain of JAK2 and tyrosines from TPOR ICD

**Page 7.** **Figure S6**. Close up of the SH2-like (SH2L) domain of human JAK2

**Page 8.** **Figure S7.** Activation loops in the TK domains of JAKs.

**Page 9.** **Figure S8.** Suggested ligand-induced activation process of the GHR-JAK2 complex

**Page 10.** **Figure S9.** Quality metrics of AFM (V2 and V3) models of the EPO-EPOR complex

**Page 11**. **Figure S10.** Quality metrics of AFM (V2) models of the GH1-GHR and PRL-PRLR complexes

**Page 12.** **Figure S11**. Quality metrics of AFM (V3) models of TPO-TPOR and CSF3-CSF3R complexes

**Page 13.** **Table S1**. Structural features and interacting partners of cytokine receptors

**Page 14.** **Table S2.** Parameters of AFM models of ligand-bound active receptor dimers, JAK2 dimers and JAK2-TMD-ICD complexes

**Page 15.** **Table S3.** Parameters of AFM models of ligand-free receptor dimers and TMD dimers

**Page 16.** **Table S4**. Interactions of hTPO with hTPOR in the AFM model of hTPO-TPOR complex

**Page 17.** **Table S5.** TM helix arrangement in partially and fully-modeled receptor complexes

**Page 18.** **Table S6:** Lipid composition of the mammalian plasma membrane

**Page 19.** **References**

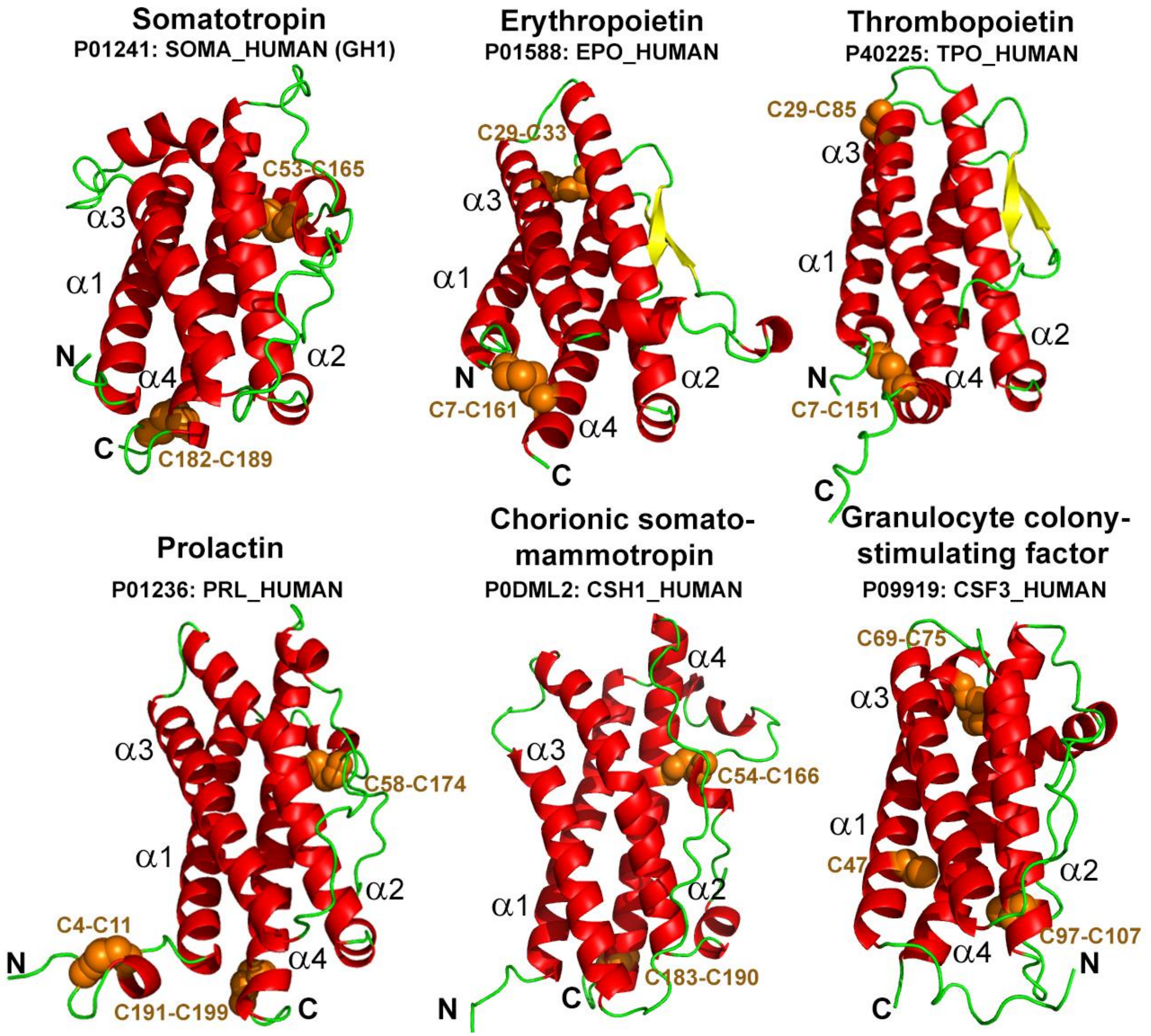


**Figure S1**. Structures of class 1 cytokines modeled by AlphaFold Multimer (AFM) in complex with their cognate receptors. The depicted cytokines are four-helical α-bundles with up-up-down-down topology and 2-3 disulfides that stabilize loop conformations. Models are shown as cartoons colored by secondary structure: red for α-helix, yellow for β-strand, green for unstructured loops. Cysteine residues and disulfides are shown by orange spheres. Residue numbers are for mature proteins (except CSF3).

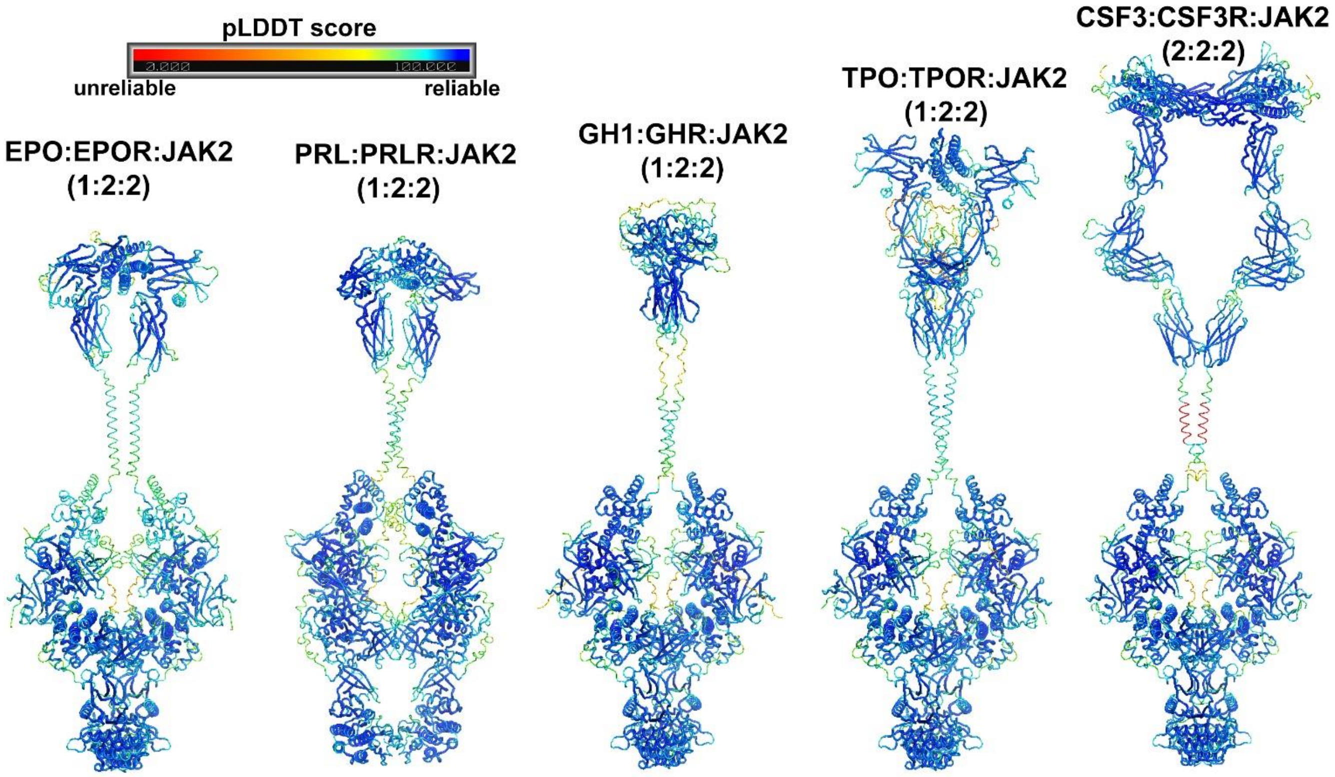


**Figure S2.** AFM-predicted models of five cytokine receptor signaling complexes colored by per residue pLDDT score.

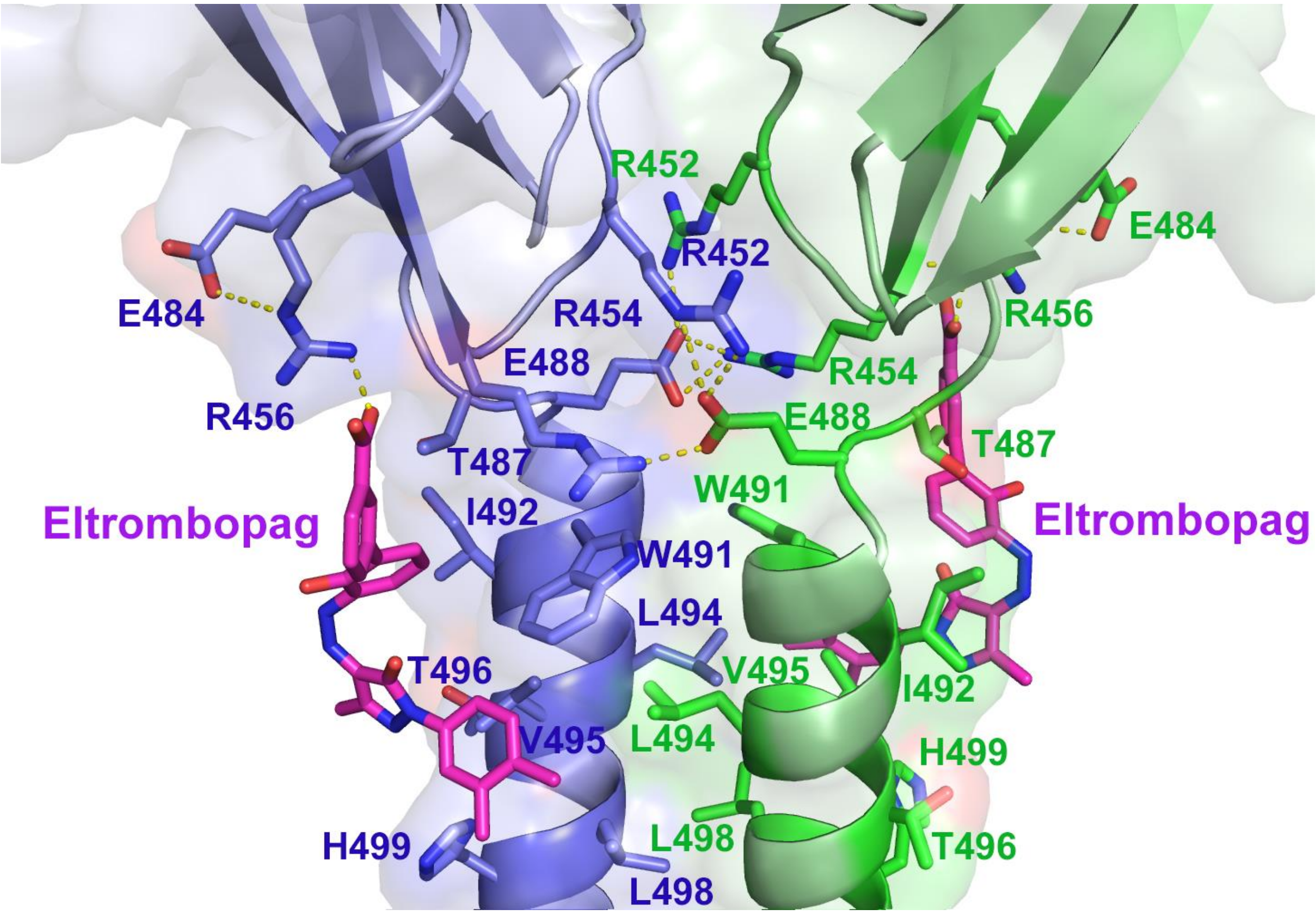


**Figure S3**. Suggested binding mode and interactions of the small molecule agonist eltrombopag with TM α-helices of the active human TPO-TPOR-JAK2 (1:2:2) signaling complex. Two eltrombopag molecules interact with the N-terminal part of the TM α-helical dimer (residues W491, I492, L494, V495, T496, L498, and H499) and R456 from the D4 domain. Neighboring residues (E488, R452, and R454) form a hydrogen bond network between D4 domains of TPOR (shown by yellow dashes). Molecules are shown by cartoon and semi-transparent surface representations colored blue and green for receptor subunits. Eltrombopag (colored magenta for Cα-atoms) and neighboring residues (colored blue and green for Cα-atoms) are shown as sticks.

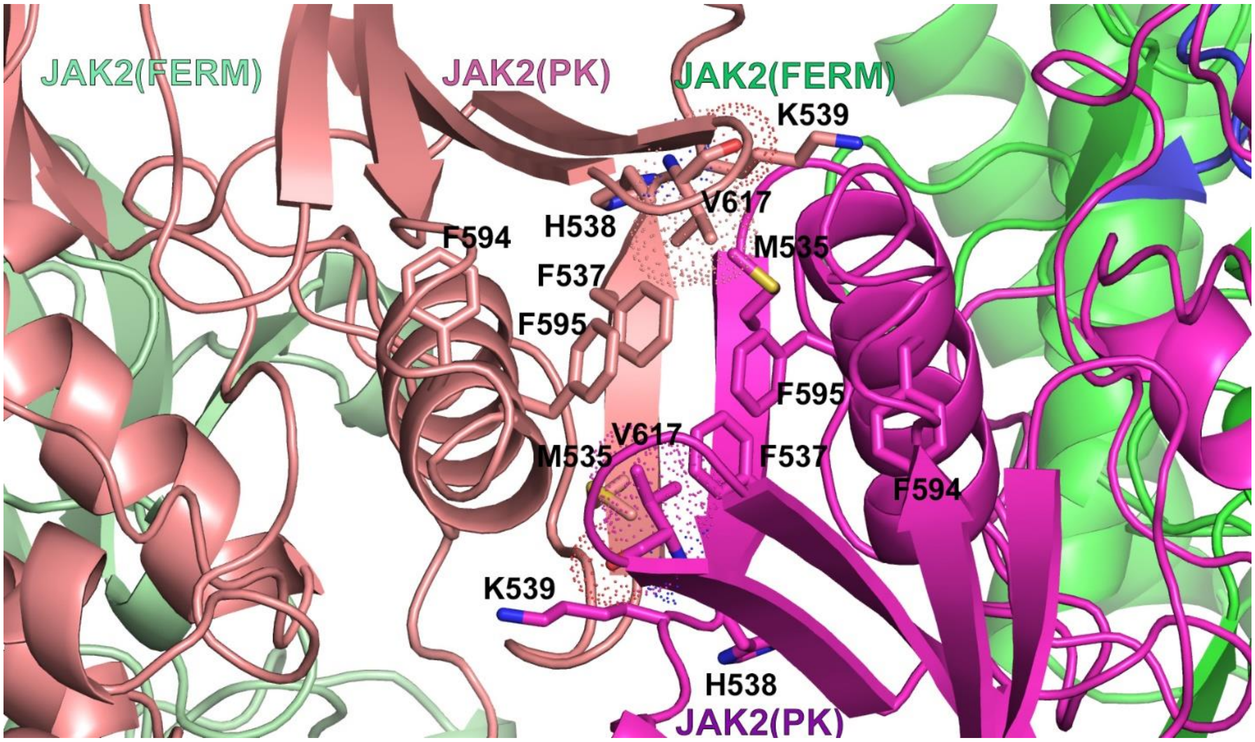


**Figure S4.** Close-up of the PK dimerization interface in the active human JAK2 dimer generated by AFM. The interface is formed by two antiparallel β-strands from PK N-lobes (cartoon representation colored pink and purple) and contains the hydrophobic cluster with V617 surrounded by aromatic residues, F537 and F595, as well as by PK residues involved in oncogenic mutations, M535, H538, K539 (shown by sticks). The oncogenic mutations (V617F, M535L, H538L, and K539L) stabilize the PK dimer by enhancing hydrophobic contacts and shape complementarity at the dimerization interface.

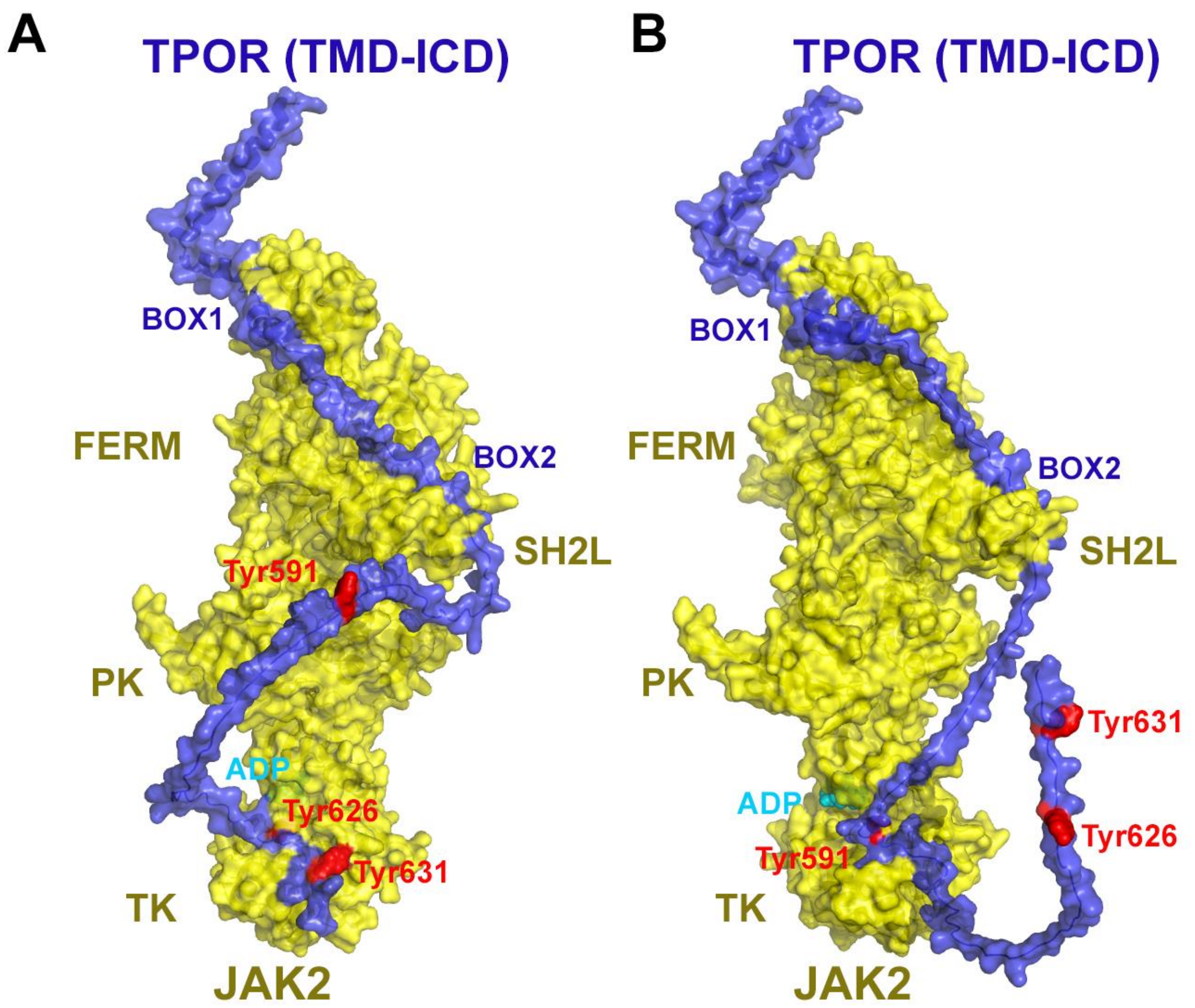


**Figure S5**. AFM modeling of interactions between TK domain of JAK2 and C-terminal tyrosine residues of TPOR that undergo phosphorylation by JAK2; Y626 (**A**) and Y591 (**B**). Due to the high flexibility of the unstructured ICD, different tyrosine residues can bind to the ligand binding pocket of the TK domain of JAK2. Protein molecules are shown using semi-transparent surfaces and cartoon representations and are colored yellow for JAK2, blue for TPOR. The ADP ligand is shown in cyan and TPOR tyrosine residues (Y591, Y626, and Y631) are colored red.

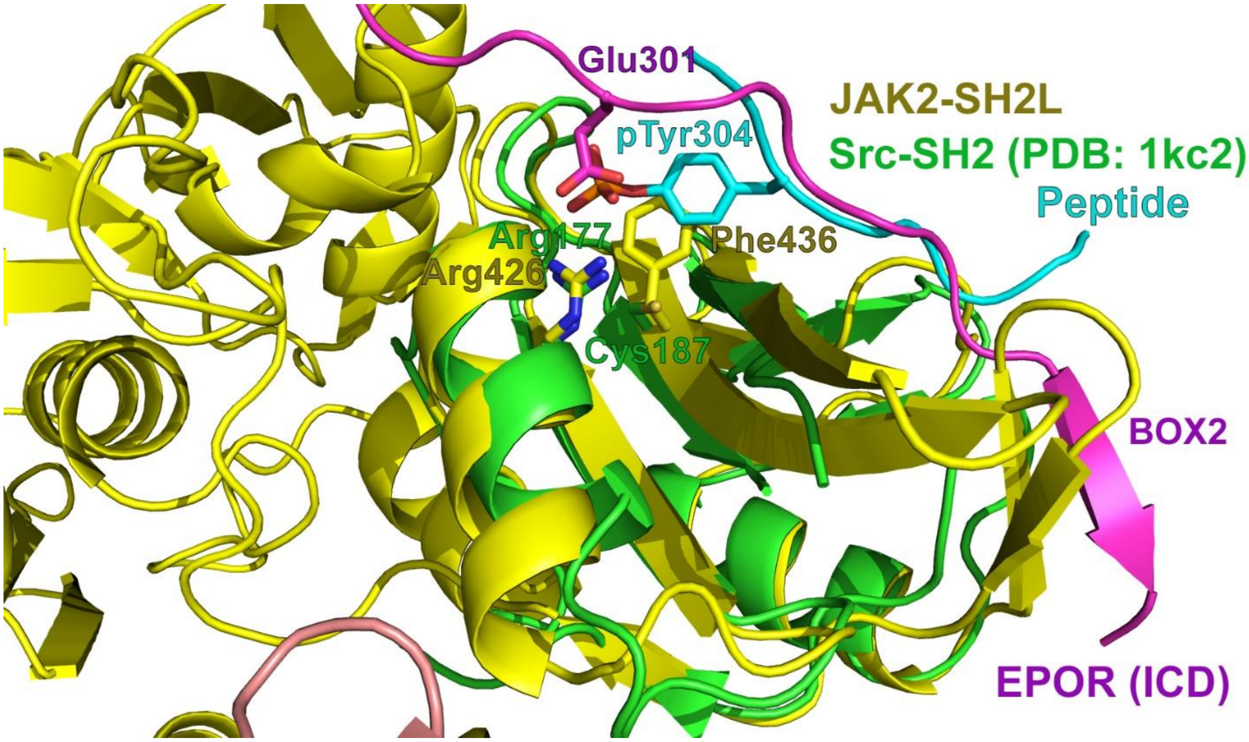


**Figure S6**. Close up of the SH2-like (SH2L) domain of human JAK2 (colored yellow) with a bound fragment of the human EPOR ICD (colored purple) from the AFM model superposed with the crystal structure of the Src-SH2 domain (colored green) with bound peptide (colored cyan) containing a phosphorylated tyrosine, pTyr304 (PDB ID: **1KC2**). SH2L domain of JAK2 has an aberrant binding site for phosphorylated tyrosine. This aberrant site carries Arg426 for binding negatively charged groups (i.e. phosphates), but lacks space for the tyrosine aromatic ring because the conserved cysteine is substituted by a bulky Phe436 residue. Therefore, the SH2L domain can specifically bind the negatively charged Glu301 from hEPOR Box 2 motif by forming ionic interactions with Arg426. Similar interactions were observed in the crystal structure of the JAK2 FERM-SH2L domain with the EPOR ICD peptide (PDB ID: **6E2Q**). It was suggested that these interactions contribute to the specificity of receptor binding [1, 2].

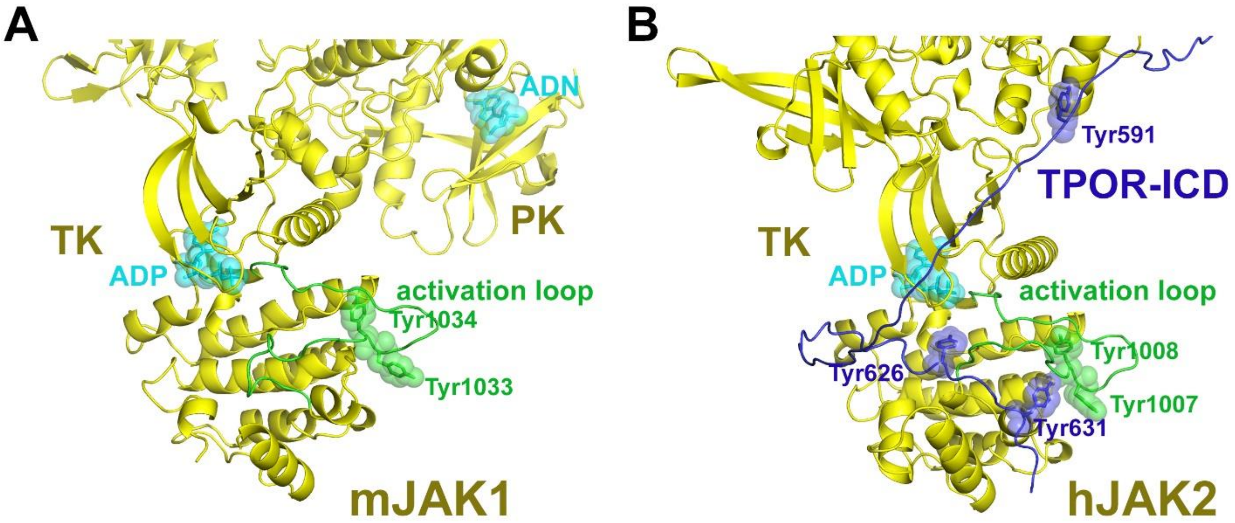


**Figure S7.** Activation loops in the TK domains of JAKs. (**A**) TK and partial PK domains from the cryo-EM-based model of mJAK1 (PDB ID: **7T6F**). (**B**) The TK domain from the AFM-generated model of hJAK2 with the fragment of TPOR ICD interacting with PK. The flexible activation loop may partially occlude the ligand binding pocket of the TK domain. Protein molecules are shown in cartoon representations colored blue for TPOR ICD, yellow for JAK2 with the activation loop of the TK domain colored green; ligands (ADP and ADN) are shown as sticks and semi-transparent cyan spheres. Tyrosine residues of TPOR, including Y626, occupying the ligand binding pocket of TK, are shown as blue sticks and semi-transparent spheres (**B**).

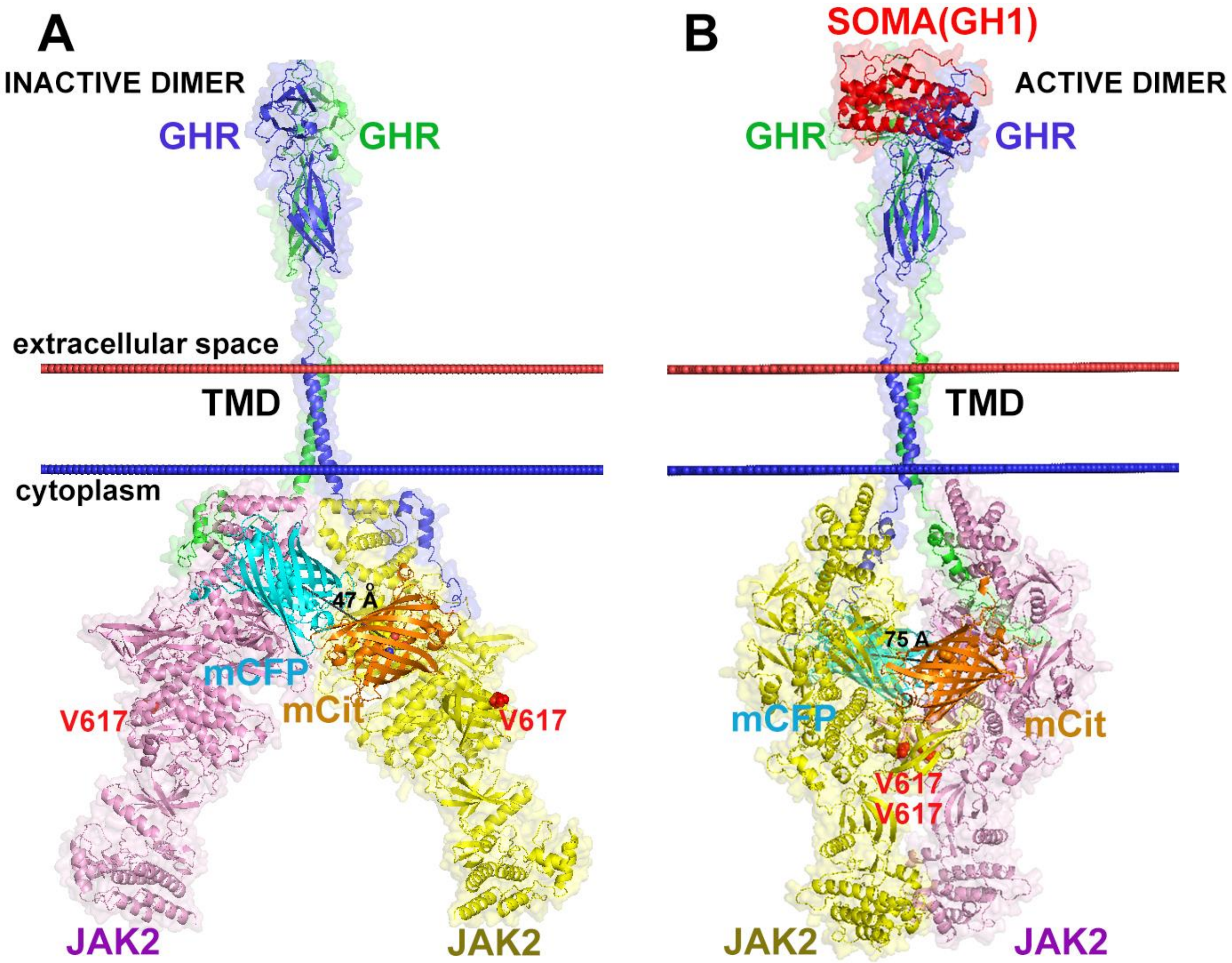


**Figure S8**. Suggested ligand-induced activation process of the GHR-JAK2 complex. AFM-generated models of ligand-free GHR-JAK2 dimer (**A**), and GHR-JAK2 dimer in the presence of GH1 ligand (**B**). Comparison with the results of FRET between GHR molecules labeled by FRET reporters (mCit and mCFP) positioned at C-terminus, 37 residues below the Box1 motif [3]. In the inactive receptor state, FRET reporters are located at the same side of the JAK2 dimer (distance between chromophores is 47 Å). However, in the active state, FRET reporters are located at the opposite sides of the JAK2 dimer (distance between chromophores is 75 Å). Protein molecules are shown as semi-transparent surfaces and cartoon representations, colored red for GH1, blue and green for GHR subunits, yellow and pink for JAK2 subunits. FRET reporters mCit and mCFP (shown as orange and cyan β-barrels, respectively) were modeled using the available CFP structure (PDB ID: **3ZFT**). V617 residue located at the dimerization interface of the PK domain is shown as red spheres.

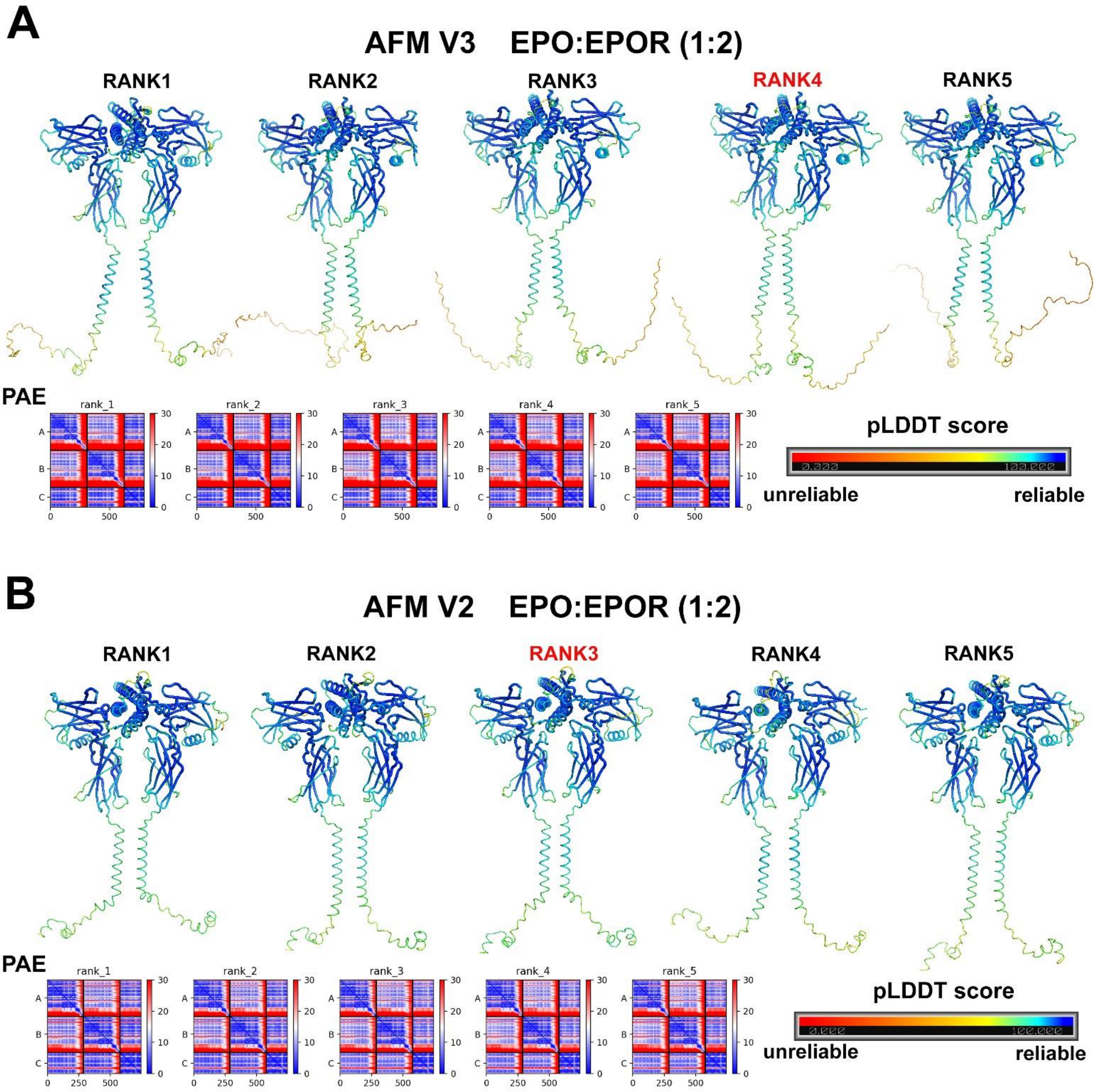


**Figure S9.** Quality metrics of AFM predicted models. (**A**) Five models predicted for EPO-EPOR (1:2) complex by AFM V3 (A) and AFM V2 (**B**). Models are colored by per residue pLDDT scores.

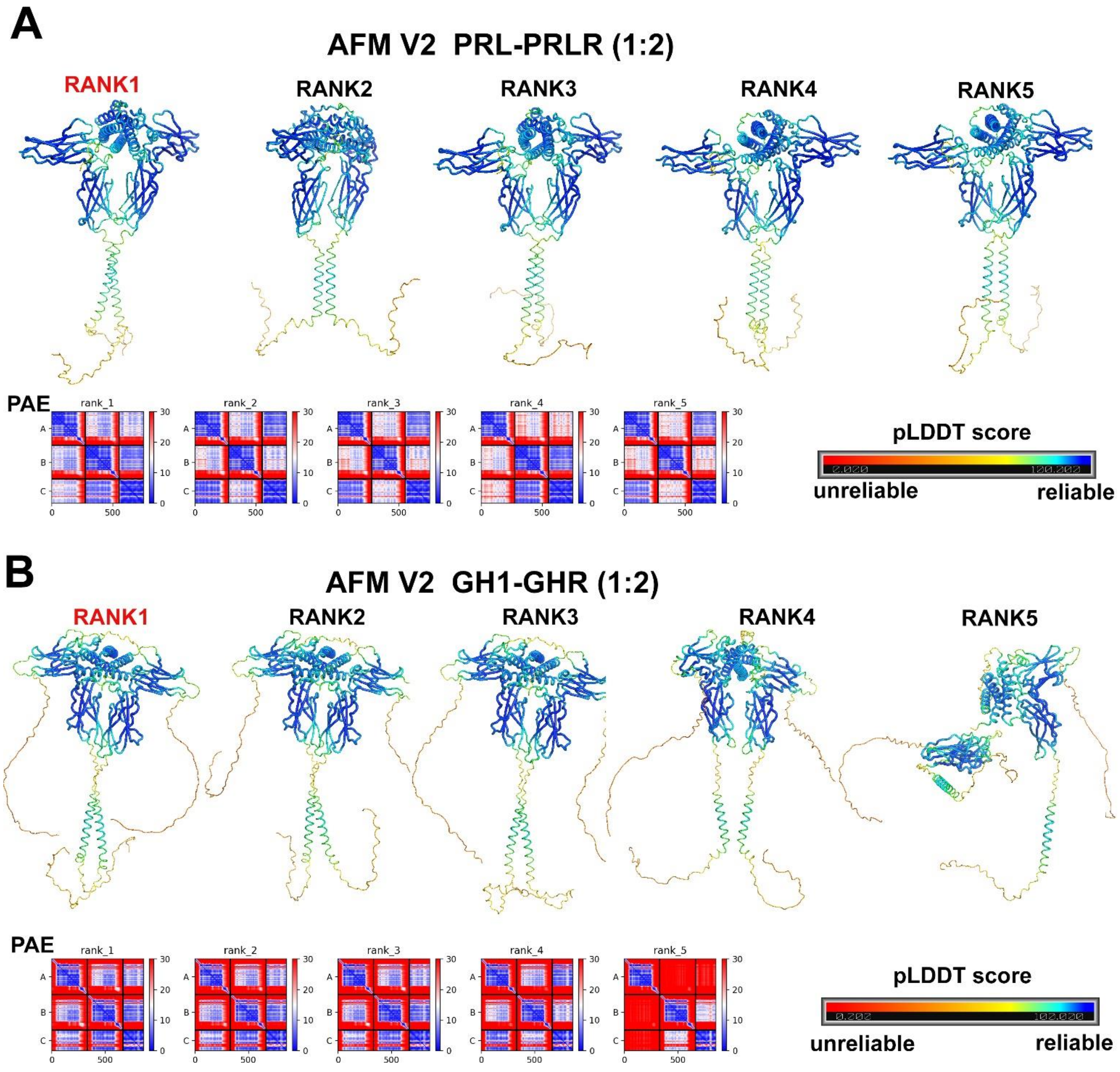


**Figure S10.** Quality metrics of AFM V2 predicted models. (**A**) Five models generated for PRL-PRLR (1:2) complex. (**B**) Five models generated for GH1-GHR (1:2) complex. Models are colored by per residue pLDDT scores.

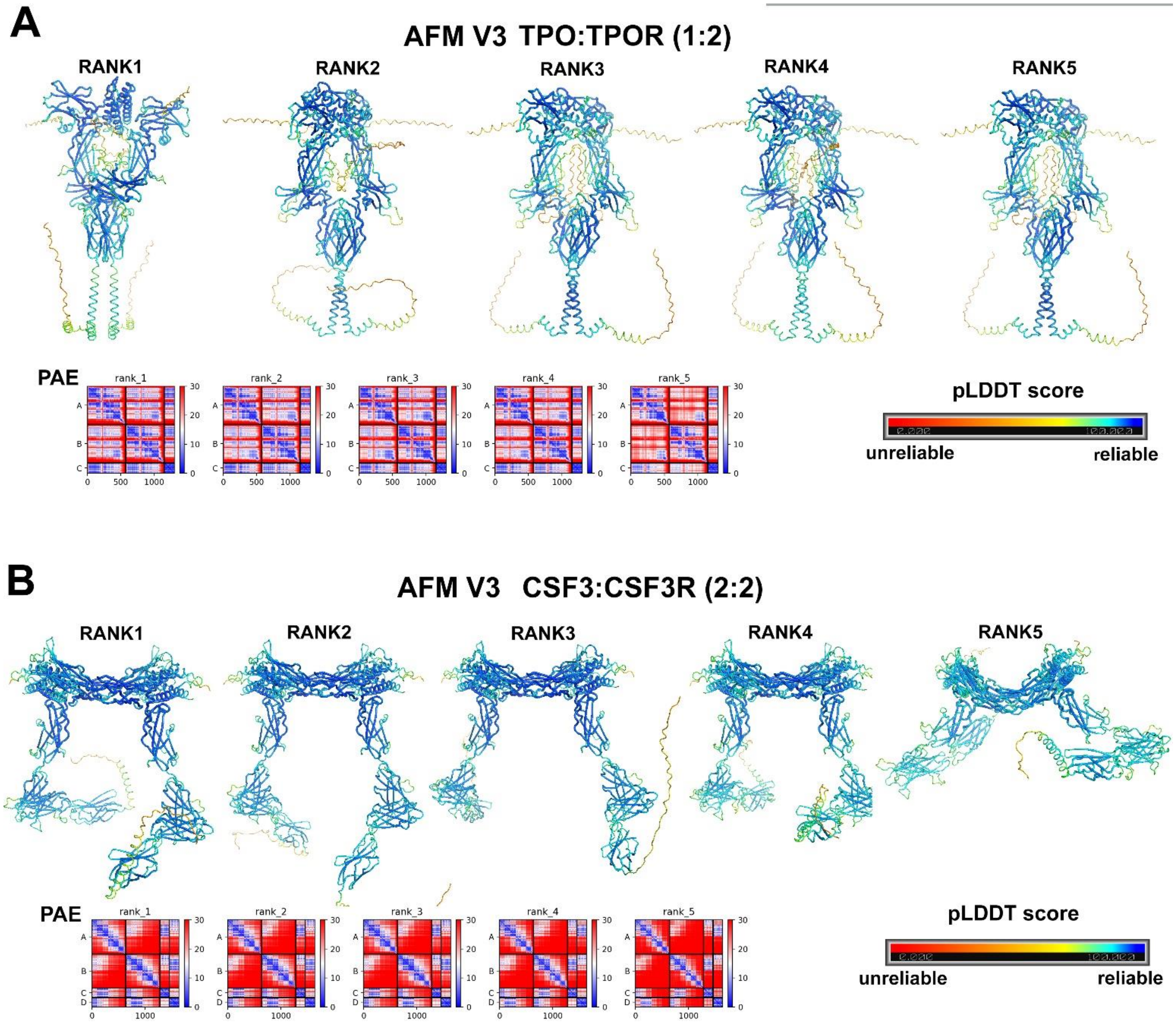


**Figure S11**. Quality metrics of AFM predicted models. (**A**) Five models predicted for TPO-TPOR (1:2) complex by AFM V3. (**B**) Five models predicted for CSF3-CSF3R (2:2) complex by AFM V3. Models are colored by per residue pLDDT scores.

**Table S1**. Structural features and interacting partners of class 1 homodimeric cytokine receptors from the JAK-STAT signaling pathway [4-6].

| **Receptor** | **Sequence length (signal peptide)** | **Disulfides** | **N-glycosylation sites** | **WSXWS motifs** | **Natural ligands** | **JAKs** | **STATs** | **SOCSs** |
|---|---|---|---|---|---|---|---|---|
| **EPOR*** | 508 (24) | 28-38, 67-83 | 52 | 209WSAWS | **EPO** | **JAK2** LYN | **STAT5A** STAT3, STAT1 | SOCS3 |
| **PRLR*** | 622 (24) | 12-22, 51-62 | 35, 80, 108 | 191WSAWS | **PRL,** CSH1, CSH2, GH1 | **JAK2** | **STAT5A,** STAT5B STAT3, STAT1 | SOCS2 |
| **GHR** | 638 (18) | 56-66, 101-112, 126-140, 259-259† | 115, 156, 200 | 240YGEFS | **GH1 (SOMA)** | **JAK2,** LYN | **STAT5B** STAT3, STAT1 | SOCS2 |
| **TPOR** | 635 (25) | 40-50, 77-93, 291-301, 334-352, 193-323#, 194-241#, 211-322# | 117, 178, 298, 358 | 269WGSWS 474WSSWS | **TPO,** CRTmut** | **JAK2,** TYK2 | **STAT5A**, STAT3, STAT1 | SOCS3 |
| **CSF3R** | 780 (24) | 26-52, 46-101, 131-142, 167-218, 177-186, 248-295, 266-309, 388-395 | 51, 93, 128, 134, 389, 474, 571, 610 | 318WSDWS | **CSF3** | **JAK1,** JAK2, JAK3 | **STAT3,** STAT5 STAT1 | SOCS3 |

* Residue numbers are for mature proteins (lacking signal peptide).
† Intermolecular disulfide
### Disulfides possibly formed between loops inside D2 and D3 domains
** CRTmut, calreticulin mutants related to MPNs
Bold characters indicate the main interacting protein.

**Table S2.** Parameters of AFM-generated models of homodimers of ligand-bound receptors, dimers of TM segments for constitutively active mutants, JAK2 dimers, and monomeric JAK2-receptor complexes. These were used as structural blocks for building complete models of active ligand-receptor-kinase signaling complexes.

| Name | Residues | pLDDT | pTM | ipTM | TM helix packing | PDB code or model | RMSD, Å |
|---|---|---|---|---|---|---|---|
| **Ligand-bound receptor homodimers** | | | | | | | |
| hEPO*-hEPOR* (1:2) | 1-166* (lig), 1-288* | 81.7 | 0.737 | 0.680 | **L+, $L^{239}$ *e*** | **1EER** | 1.6 (560/592) |
| mEPO*-mEPOR (1:2) | 1-166* (lig) 1-314 | 78.5 | 0.687 | 0.645 | **L+, $S^{238}$ e** | **1EER** | 1.06 (462/592) |
| hGH1*-hGHR (1:2) | 1-191* (lig) 1-332 | 70.3 | 0.648 | 0.639 | **L+, $F^{273}$ *d-*** | **3HHR 5OEK** | 1.2 (549/573) 3.1 (48/48) |
| hPRL*-hPRLR* (1:2) | 1-199* (lig) 1-276* | 80.2 | 0.680 | 0.635 | **R-, $AxxxA^{222}xxxL$** | **3NPZ** | 1.8 (536/587) |
| hCSH1*-hPRLR* (1:2) | 1-191* (lig) 1-276* | 80.8 | 0.665 | 0.584 | **R-, $LxxxW^{230}xxxL$** | **1F6F** | 1.85 (403/698) |
| hGH1*-hPRLR* (1:2) | 1-191* (lig) 1-276* | 82.2 | 0.713 | 0.641 | **R-, $CxxxV^{229}xxxA$** | **1BP3** | 1.1 (322/383) |
| hTPO*-hTPOR (1:2)# | 1-153* (lig) 1-635 | 75.1 | 0.67 | 0.639 | **L+, $S^{505}$ *a*, $H^{499}$ *b*** | None | |
| hCSF3-hCSF2R (1:1) | 30-207 (lig) 1-667 | 83.1 | 0.664 | 0.788 | N/A | **2D2Q** | 1.0 (409/466) |
| **JAK2 homodimer** | | | | | | | |
| hJAK2 dimer | 1-1132 | 79.5 | 0.743 | 0.672 | 224-224 distance 29.3 Å | **7T6F 8EWY** | 2.5 (1898/2164) 3.0 (2052/2172) |
| **JAK2-receptor complexes (1:1)** | | | | | | | |
| hJAK2-hEPOR* (D2-TM-ICD) | 1-1132, 120-372* | 82.9 | 0.769 | 0.631 | L+ | **7T6F** (FERM-SH2L-PK) | 2.48 (618/762) |
| hJAK2-hGHR (D2-TM-ICD) | 1-1132, 148-390 | 82.5 | 0.754 | 0.634 | L+ | **7T6F** (FERM-SH2L-PK) | 2.47 (620/762) |
| hJAK2-hPRLR* (TM-ICD) | 1-1132, 205-295* | 85.6 | 0.838 | 0.727 | R- | **7T6F** (FERM-SH2L-PK) | 2.45 (620/762) |
| hJAK2-hTPOR (TM-ICD) | 1-1132, 488-635 | 82.8 | 0.808 | 0.552 | L+ | **7T6F** (FERM-SH2L-PK) | 2.63 (636/762) |
| hJAK2-hCSF3R (TM-ICD) | 1-1132, 606-708 | 84.6 | 0.826 | 0.612 | L+ | **7T6F** (FERM-SH2L-PK) | 2.58 (621/762) |

* Residue number are for mature proteins.

The AFM parameters (pLDDT, pTM, ipTM) are provided for a single model selected for further modeling and analysis out of 5 models generated by AFM.

Type of helix arrangement in dimers (L+ or R-) as defined by the sign of the crossing angle: L+, left-handed (coiled coil) dimer with a positive crossing angle and heptad repeat, or R-, right-handed dimer with a negative crossing angle and the tetrad (i.e. GxxxG) repeat motif. Letters for left-handed dimers indicate the position of a reference residue in the $(abcdefg)_n$ heptad repeat motif (*a* and *d* positions are located at the helix-helix interface). RMSD column includes the number of overlapped residues in the structural superposition divided by total number of residues in the structure (in parentheses).

Bold characters indicate a TM helix packing consistent with the final structure of the active receptor dimer.

### calculated by AFM V3. Other models were calculated by AFM V2.

**Table S3.** Parameters of AFM-generated models of ligand-free receptor dimers and dimers of TM α-helical segments.

| Name | Residues | pLDDT | pTM | ipTM | TM helix packing | PDB entry or another model | RMSD, Å* |
|---|---|---|---|---|---|---|---|
| **Ligand-free receptor homodimers** | | | | | | | |
| hEPOR* # | 1-271*<br>1-269* | 76.8 | 0.456 | 0.154 | L+, L239 *b* | 1eer | 8.94 (412/538) |
| mEPOR* | 1-190* | 75.7 | 0.496 | 0.258 | L+, $S^{238}$ *a* | 1eer | 5.6 (408/580) |
| hGHR | 1-310 | 70.8 | 0.503 | 0.352 | L+, $F^{273}$ *e* | 3hhr | 7.5 (384/573) |
| hPRLR* | 2-279* | 82.4 | 0.474 | 0.163 | L+, $A^{222}$ *a* | 3npz | 8.6 (391/480) |
| hTPOR# | 26-550<br>1-552 | 72.4 | 0.526 | 0.444 | R- $H^{499}$ out | Final model | 4.8 (952/1104) |
| hCSF3R (D5-D6-TM segment) | 421-660 | 79.2 | 0.389 | 0.120 | R-, $T^{640}$ out | Final model | 8.1 (434/480) |
| **Homodimers of TM α-helices*** | | | | | | | |
| hEPOR* | 209-288* | 54.6 | 0.393 | 0.337 | L+, $L^{239}$ *a* | Final model<br>TMDOCK | 2.9 (57/58)<br>1.7 (53/58) |
| mEPOR* | 208-287* | 52.8 | 0.364 | 0.306 | L+, $S^{238}$ *a* | Final model<br>TMDOCK | 3.1 (56/58)<br>2.0 (56/58) |
| hGHR | 240-325 | 52.0 | 0.394 | 0.356 | L+, $F^{273}$ *a* | 5oek<br>Final model | 3.6 (53/54)<br>1.6 (51/54) |
| hPRLR* | 191-249* | 62.7 | 0.516 | 0.467 | **R-, AxxxA$^{222}$xxxL** | Final model<br>TMDOCK | 2.7 (68/68)<br>2.2 (48/50) |
| hPRLR* CAM (∆10-186) | 1-9+187-276 | 49.7 | 0.384 | 0.360 | **R-, AxxxA$^{222}$xxxL** | Final model<br>TMDOCK | 1.9(67/68)<br>3.3(49/50) |
| hPRLR* CAM (∆1-210) | 211-276 | 62.0 | 0.492 | 0.461 | **R-, AxxxA$^{222}$xxxL** | Final model<br>TMDOCK | 1.7 (60/68)<br>3.6 (50/50) |
| hTPOR CAM H499L/W515K | 488-549 | 71.9 | 0.463 | 0.373 | L+, $S^{505}$ *d*, $H^{499}$ *e* | Final model<br>TMDOCK | 1.2 (52/54)<br>1.4 (48/58) |
| hTPOR CAM (L498W/H499Y) | 488-549 | 73.5 | 0.49 | 0.404 | L+, $S^{505}$ *d*, $H^{499}$ *e* | Final model<br>TMDOCK | 1.2 (52/54)<br>1.4 (48/58) |
| mTPOR | 456-533 | 63.6 | 0.47 | 0.397 | L+, $S^{498}$ *d*, $L^{492}$ *e* | Final model<br>TMDOCK | 2.8 (54/54)<br>1.9 (56/58) |
| hCSF3R | 606-669 | 60.1 | 0.483 | 0.442 | **L+, $T^{640}$ *a*** | Final model<br>TMDOCK | 0.8 (49/54)<br>0.8 (41/50) |
| hCSF3R CAM (T640N) | 606-669 | 58.8 | 0.439 | 0.389 | **L+, $T^{640}$ *a*** | Final model<br>TMDOCK | 0.7 (48/54)<br>0.9 (44/50) |

See legend for Table S2. Bold characters indicate TM helix arrangement similar to that in the models of final ligand-receptor-kinase signaling complexes. CAM, constitutively active mutants.
### calculated by AFM V3. Other models were calculated by AFM V2.

**Table S4**. Interactions of hTPO with hTPOR in the AFM-generated model of the ligand-bound hTPOR dimer.*

| Site 1 | | Site2 | |
|---|---|---|---|
| TPO* | **TPOR** | **TPO*** | **TPOR** |
| **L16** | I263 | **R10** | **E261** |
| **D45** | R102 | V11 | **F104** |
| **F46** | **L103, F104** | **K14** | E160 |
| **L48** | **F45, L103, L265** | **R17** | D163 |
| K52 | E46 | **R98** | E99 |
| H133 | F126 | L99 | **L103, F104** |
| K136 | D128 | L101 | F105 |
| R140 | **D261** | | |
| F141 | **F104**, F164, **L265** | | |
| L144 | F164 | | |

* Residue numbers are for mature protein (TPO).

Bold characters indicate hTPOR residues (F45, L103, F104, D261, and L265) that are involved in hTPO binding, in accordance with mutagenesis studies [7, 8] and hTPO residues (R10, K14, R17, K52, R98, H133, K136, F141, L144) that are essential for binding to hTPOR [9, 10].

**Table S5.** TM helix arrangement in final models of complete ligand-receptor-kinase complexes, in models of ligand-bound and ligand-free receptor dimers, and in dimers of isolated TM segments calculated by AFM [11] and TMDOCK [12].

| Name | Complete complex | Ligand-bound receptor dimer | Ligand-free receptor dimer | TM helix dimer (AFM) | TM helix dimer (TMDOCK) |
|---|---|---|---|---|---|
| **hEPOR*** | L+, $L^{239}$ *e* | L+, $L^{239}$ *e* | R- | L+, $L^{239}$ *a* | no association |
| **mEPOR*** | L+, $S^{238}$ *e* | L+, $S^{238}$ *e* | L+, $S^{238}$ *a* | L+, $S^{238}$ *a* | L+, $S^{238}$ *a* |
| **hGHR** | L+, $F^{273}$ *d* | L+, $F^{273}$ *d* | L+, $F^{273}$ *e* | L+, $F^{273}$ *a* | L+, $F^{273}$ *c* |
| **hPRLR*** | R-, AxxxA$^{222}$xxxL | R-, AxxxA$^{222}$xxxL | L+ $A^{222}$ *a* | R-, AxxxA$^{222}$xxxL (WT,CAM**) | R-, SxxxC$^{225}$xxxV |
| **hCSF3R** | L+, $T^{640}$ *a* | N/A | N/A | L+, $T^{640}$ *a* (WT,CAM**) | L+, $T^{640}$ *a* |
| **hTPOR** | L+, $S^{505}$ *a*, $H^{499}$ *b* | L+, $S^{505}$ *a*, $H^{499}$ *b* | R- | L+, $S^{505}$ *d*, $H^{499}$ *e* (CAM)** | L+, $S^{505}$ *d*, $H^{499}$ *e* |

Cells marked by gray indicate helix arrangements similar to that in the final ligand-receptor-kinase complexes.
Type of helix arrangement in dimers (L+ or R-) is defined by the sign of the crossing angle: L+, left handed dimer (coiled coil) with positive crossing angle and the $(abcdefg)_n$ heptad repeat motif, and R-, right-handed dimer with a negative crossing angle and the tetrad (i.e. GxxxG) repeat motif. Letters for left-handed dimers (bold character) indicate the position of a reference residue in the heptad repeat motif, where *a* and *d* positions form the dimerization interface.
* Residue numbers are for mature proteins.
** Calculated for TMDs of constitutively active mutants (CAM): hPRLR (Δ10-186, Δ1-210), hCSF3R (T640N), and TPOR (H499L/W515K and L498W/H499Y).

**Table S6:** Lipid composition of the mammalian plasma membrane (number of specified lipid molecules in each leaflet).

| Lipid name | Lipid Head / Tail | GHR | | CSF3R, EPOR, PRLR, TPOR | |
|---|---|---|---|---|---|
| | | Outer | Inner | Outer | Inner |
| **POPC** | PC(16:0/18:1(9Z)) | 64 | 28 | 48 | 21 |
| **PLPC** | PC(16:0/18:2(9Z,12Z)) | 88 | 44 | 66 | 33 |
| **PAPE** | PE(16:0/20:4(5Z,8Z,11Z,14Z)) | 12 | 48 | 9 | 36 |
| **POPE** | PE(16:0/18:1(9Z)) | 12 | 56 | 9 | 42 |
| **POPI** | PI(16:0/18:1(9Z)) | 0 | 20 | 0 | 15 |
| **PAPS** | PS(16:0/20:4(5Z,8Z,11Z,14Z)) | 0 | 44 | 0 | 33 |
| **POPA** | PA(16:0/18:1(9Z)) | 0 | 4 | 0 | 3 |
| **SSM** | SM(d18:1/18:0) | 44 | 20 | 33 | 15 |
| **NSM** | SM(d18:1/24:1) | 44 | 20 | 33 | 15 |
| **CMH** | GlcCer(d18:1/16:0) | 16 | 0 | 12 | 0 |
| **CHOL** | Cholesterol | 148 | 116 | 111 | 87 |
| **Total** | | 428 | 400 | 321 | 300 |

Lipids head groups: PC, phosphatidylcholine; PE, phosphatidylethanolamine; PI, phosphatidylinositol; PS, phosphatidylserine; SM, sphingomyelin; GlcCer, glucosylceramide.
